# The histone variant macroH2A1.1 regulates RNA Polymerase II paused genes within defined chromatin interaction landscapes

**DOI:** 10.1101/2020.01.29.924704

**Authors:** Ludmila Recoules, Alexandre Heurteau, Flavien Raynal, Nezih Karasu, Fatima Moutahir, Fabienne Bejjani, Isabelle Jariel-Encontre, Olivier Cuvier, Thomas Sexton, Anne-Claire Lavigne, Kerstin Bystricky

## Abstract

The histone variant macroH2A1.1 (mH2A1.1) plays a role in cancer development and metastasis-related processes. To determine the underlying molecular mechanisms, we mapped genome-wide localization of endogenous mH2A1.1 in the human breast cancer cell MDA-MB 231. We demonstrate that mH2A1.1 specifically binds to active promoters and enhancers in addition to facultative heterochromatin. Selective knock-down of mH2A1.1 deregulates expression of hundreds of highly active genes. Depending on the chromatin landscape, mH2A1.1 acts through two distinct molecular mechanisms. The first is to limit excessive transcription in a predefined environment and relies on domain recruitment of mH2A1.1 at the promoter and gene body. The second mechanism is specific to RNA Pol II (Pol II) paused genes. It requires recruitment of mH2A1.1 restricted to the TSS of these genes. Moreover, we show that these processes occur in a predefined local 3D genome organization and are largely independent of enhancer-promoter looping. Among the genes activated by mH2A1.1, genes regulating mammary tumor cell migration are mostly dependent on Pol II release for their expression level, unlike other categories of mH2A1.1-regulated genes. We thus identified an intriguing new mode of transcriptional regulation by mH2A1.1 and propose that mH2A1.1 serves as a transcriptional modulator with a potential role in assisting the conversion of promoter-locked RNA polymerase II into a productive and elongated Pol II.

## Introduction

Histone post-translational modifications, DNA-binding factors and architectural proteins regulate genome organization and dynamics (Luger et al., 2012; Venkatesh & Workman, 2015). In addition, histone variants replace canonical histones in a locus-specific manner, which endows chromatin with properties required to fine-tune DNA accessibility and functions (Buschbeck & Hake, 2017).

Among the histone variants, macroH2A1 (mH2A1), a vertebrate-specific (Pehrson & Fuji, 1998; Rivera-Casas et al., 2016) histone H2A variant, is composed of an N-terminal “H2A- like” domain (64 % identical to H2A) and a C-terminal 25 kDa “macro” domain. These two domains are joined by an unstructured 41 amino acid long “linker” domain that positions the macro domain outside of the nucleosome (Gamble & Kraus, 2010). Expression of the highly conserved *H2AFY* gene produces two splicing isoforms, mH2A1.1 and mH2A1.2, whose sequences differ in a 30 amino-acid region within the macro domain (Gamble & Kraus, 2010).

mH2A1 was originally found to be enriched on the transcriptionally silent X chromosome (Costanzi & Pehrson, 1998). mH2A1 is also present at autosomes, forming large domains in association with histone marks associated with heterochromatin, such as H3K27me3 and H3K9me3 (Douet et al., 2017; Gamble et al., 2010; Sun et al., 2018). *In vitro* studies have demonstrated that nucleosomal mH2A1 interferes with binding of the transcription factor NF-kB, inhibits nucleosome sliding by the remodeling complex SWI/SNF and initiation of RNA polymerase II (Pol II) transcription (Angelov et al., 2003; Doyen et al., 2006). Therefore, mH2A1 is generally believed to play a role in transcriptional repression. However, in many cases, the presence of mH2A1 correlates also with active transcription of a subset of genes (Changolkar et al., 2010; H. Chen et al., 2014; Dell’Orso et al., 2016; Gamble et al., 2010; Podrini et al., 2015; Wan et al., 2017). Thus, the roles of mH2A1 in regulating gene expression are seemingly contradictory and the underlying mechanisms are still not well characterized.

The two mH2A1 splice variants exhibit tissue- and cell-specific expression patterns (Posavec Marjanović et al., 2017). In normal cells, the mH2A1.2 isoform appears ubiquitously expressed (Cantariño et al., 2013; J C Sporn et al., 2009; Judith C Sporn & Jung, 2012) while mH2A1.1 is only expressed in differentiated cells with low proliferation rates (Cantariño et al., 2013; J C Sporn et al., 2009; Judith C Sporn & Jung, 2012). Notably, the mH2A1.1 “macro” domain can bind NAD^+^ metabolites (Kustatscher et al., 2005) and interact with the DNA-damage repair and chromatin remodeling factor PARP1 (Poly-(ADP-Ribose) Polymerase 1) (H. Chen et al., 2014; Marjanović et al., 2018; Ouararhni et al., 2006; Ray Chaudhuri & Nussenzweig, 2017). Interaction between mH2A1.1 and PARP1 was shown to be important during DNA damage, stress responses (Kim et al., 2018; Xu et al., 2012), mitochondrial respiratory capacity (Marjanović et al., 2018) and transcription (H. Chen et al., 2014; Ouararhni et al., 2006) in fibroblast and epithelial cancerous cells. In tumors, expression of the mH2A1.1 isoform is frequently reduced compared to normal tissues, suggesting that this isoform is a tumor suppressor (Cantariño et al., 2013; A.-C. Lavigne et al., 2014; Judith C Sporn & Jung, 2012). Interestingly, in immortalized human mammary epithelial cells, mH2A1interferes with the epithelial-mesenchymal transition (EMT), and its reciprocal, the mesenchymal-epithelial transition (MET) processes required for metastases development in a tumor context (Hodge et al., 2018; Pliatska et al., 2018). However, in highly metastatic cancers such as triple-negative breast cancers (TNBCs), increased expression levels of mH2A1.1 correlate with poor prognosis (A.-C. Lavigne et al., 2014). The role of macroH2A1.1 in controlling the properties of tumor cells could be dependent on cellular context and remains to be clarified.

In this work, we identified and characterized the role of mH2A1.1 in the regulation of gene expression in TNBC cells. We found that mH2A1.1 modulates the expression of hundreds of highly expressed genes, while mH2A1.1 deficiency does not affect the expression of silent or low expressed genes. Many of these mH2A1.1-regulated genes are involved in cytoskeletal organization, certainly giving mH2A1.1 its role in controlling the migratory properties of these tumor cells. This transcriptional function of mH2A1.1 is however bifunctional, with inhibitory or stimulatory of target gene. Although not requiring ad-hoc rewiring of promoter-enhancer contacts, this functional dichotomy clearly depends on the chromatin landscape in which these genes are located and relies on differential recruitment of mH2A1.1. The activating effect of mH2A1.1 requires tight recruitment of mH2A1.1 to the TSS of related genes. Conversely, genes inhibited by mH2A1.1 recruit this histone variant over larger domains, present further upstream and downstream of the TSS. Mechanistically, we determined that the expression level of mH2A1.1-activated genes is dependent on the Pol II pause process. Deficiency in mH2A1.1 affects Pol II turnover at their TSS, potentially by inhibiting the release of paused Pol II. Our work identifies and clarifies for the first time the ambivalence of the transcriptional functions of mH2A.1.1 in cancer cells, linking its recruitment type to its mode of action.

## Results

### mH2A1.1 regulates expression of hundreds of genes

In order to characterize the function of mH2A1.1 in triple negative breast cancer (TNBC), we performed RNA-seq data in the MDA-MB231 cell line which expresses mH2A1.1 at higher levels than other types of breast cancer cell lines (**S1A, B Fig**) (A.-C. Lavigne et al., 2014). We compared gene expression levels between WT cells and cells in which the mH2A1.1 isoform but not mH2A1.2 protein expression was abolished by RNAi (KD, **Fig. 1A ; S1C-E Fig**). Among the 945 genes whose expression was significantly modified in the mH2A1.1 KD cells, 533 genes (56.3%) were down-regulated (called mH2A1.1-activated genes or AG) and 412 genes (43.7%) were up-regulated (called mH2A1.1-repressed genes or RG) (**Fig 1A, S1 Table**). In general, gene expression changes induced by mH2A1.1 depletion are moderate (**Fig 1A**). Altered gene expression was confirmed by RT-qPCR on a subset of genes using two different siRNAs directed against mH2A1.1 (**S1F-H Fig**). All mH2A1.1-regulated genes, both RG and AG, were found among the moderately to highly expressed genes in WT MBA-MB231 cells (**Fig 1B, C**). Silenced genes in MDA-MB231 cells were not activated upon mH2A1.1 depletion (**Fig 1B, C**). We concluded that mH2A1.1 participates in fine-tuning actively transcribed genes expression. We next asked if the role of mH2A1.1 variant in controlling expression of active genes depends on its association with certain genomic regions including gene regulatory regions and specific epigenetic contexts.

**Fig 1.**
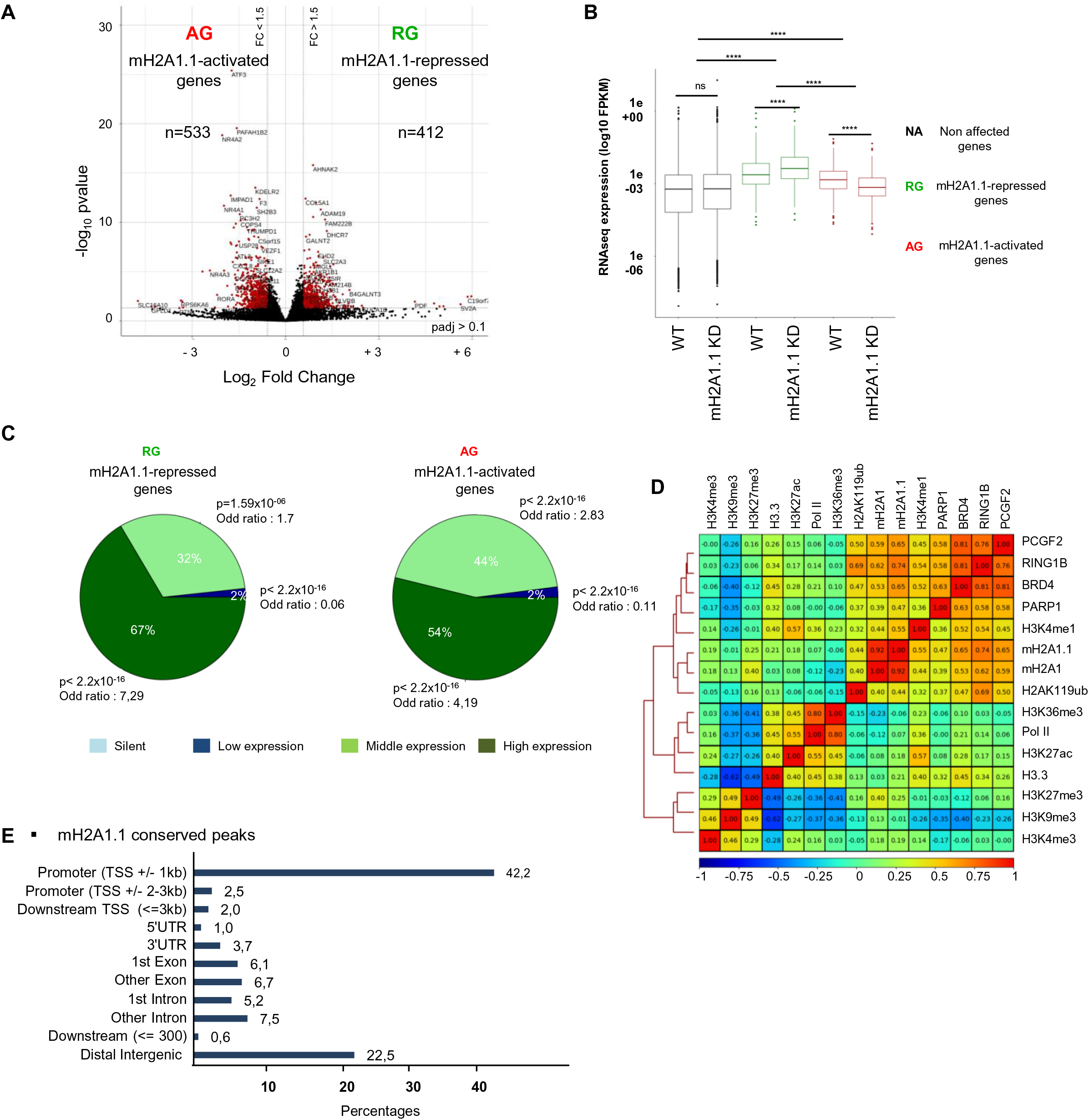
The histone variant mH2A1.1 regulates the expression of hundreds of genes in MDA-MB 231 cells. (A) Volcano plot showing fold change of gene expression in mH2A1.1 KD compared to WT MDA-MB231 cells. Red dots represent significantly deregulated genes with a fold change > 1.5 and p-adj < 0.1. (B) Boxplot comparing gene expression (FPKM) of indicated genes between control (WT) and mH2A1.1 KD conditions. **** = p-value < 2.2x10^-16^; ns = non-significant. (C) Pie charts showing proportion of mH2A1.1 regulated-genes in 4 groups categorized by gene expression levels in control cells, as indicated. Enrichment of mH2A1.1-target genes with categories of genes are measured using fisher exact tests. p-values (p) and the Odd ratios are shown. (D) Whole genome spearman correlation heatmap of mH2A1.1, mH2A1 and a series of histone modifications and chromatin associated factor ChIP-seq data, as indicated. Pearson coefficient correlations (PCC, r) are given. Red and blue colours denote high correlation (r close to 1) and anti-correlation (r close to -1), respectively. (E) Proportions of different genomic features associated with mH2A1.1 conserved peaks. mH2A1.1 “conserved” peaks correspond to the common peaks between mH2A1.1 specific ChIP-seq and mH2A1 ChIP-seq and are used for further analysis.

### mH2A1.1 associates with gene regulatory regions

We developed a ChIP-grade polyclonal antibody that exclusively recognizes mH2A1.1 (Ab αmH2A1.1) (**S2 Table and S2A-D Fig**) and generated ChIP-seq data of mH2A1.1 (**S3 Table**). The obtained dataset was compared to the one obtained using a commercially available ChIP-grade antibody (Ab37264, Ab αmH2A1) directed against total mH2A1 (**S2, S3 Tables and S2E-G Fig**). The two datasets were highly similar with a Pearson coefficient correlation (PCC) of 0.92 (**Fig 1D**). Therefore, we decided to conserved common peaks between the two ChIP-seq data for further analysis (Materials and Methods). We identified 11.849 mH2A1.1 peaks, covering ≈ 7 % of the genome. Analysis of the genomic distribution of mH2A1.1 showed that the vast majority of mH2A1.1 peaks correspond to annotated promoters (TSS +/- 1 kb), while 22% of mH2A1.1 peaks were associated with distal intergenic regions (**Fig 1E**). We confirmed the enrichment of mH2A1.1 in a subset of regions corresponding to ChIP-seq peaks by ChIP-qPCR in WT cells, as well as its decrease in mH2A1.1 KD cells (two mH2A1.1-specific RNAi), using either mH2A1.1 or mH2A1-specific antibodies (**S3 Fig)**.

### mH2A1.1 binds promoters of its target genes

We next asked if mH2A1.1 binding occurred in specific chromatin environments. Genome-wide, we analyzed the correlation between mH2A1.1 and a subset of heterochromatin marks (H3K9me3, H3K27me3, H2AK119ub), chromatin-bound components (Pol II, BRD4, RING1B, PARP1, PCGF2) as well as euchromatin marks (H3K4me1, H3K4me3, H3K36me3, H3K27ac, H3.3) (**Fig 1D**). mH2A1.1 positively correlated with its well documented partner PARP1 in the MDA-MB231 cell line (**Fig 1D**) (PCC of 0.47) (H. Chen et al., 2014). As expected, mH2A1.1 (but mainly total mH2A1) also associated with broad H3K27me3-marked chromatin domains (PCC of 0.25 for mH2A1.1 & 0.40 for mH2A1) (**Fig 1D, S4A-B Fig**). In this TNBC cell line, H3K9me3 is known to be overrepresented (Segal et al., 2018; Yokoyama et al., 2013), explaining its largely overlaps with mH2A1.1 binding at H3K27me3-marked domains (**S4A Fig**). Interestingly, we found that in heterochromatin domains comprising both H3K9me3 and H3K27me3, the level of the two histone marks tended to be inversely proportional (**S4B-C Fig**). Moreover, we found that high H3K27me3-H3K9me3 difference is mainly associated with genomic regions whereas low H3K27me3-H3K9me3 difference is more associated with intergenomic regions (**S4D Fig**). We propose that in this cell line, in which H3K9me3 is over-present, difference in the intensity signal between H3K27me3 and H3K9me3 could be used to distinguish “facultative-like” heterochromatin and “constitutive-like” heterochromatin. To decipher whether mH2A1.1 preferentially overlaps with H3K9me3- or H3K27me3-marked heterochromatin domains, we analyzed its association depending on the difference in the signal intensity of both marks. We found that mH2A1.1 (and PARP1) binding were proportional to the abundance of H3K27me3 minus H3K9me3 (**S4B, S4E, S4F Fig**), indicating that these two proteins predominantly associate with “facultative-like heterochromatin” in this TNBC breast cancer cell.

When specifically examining the chromatin landscape at promoters (TSS +/- 1kb), we found that mH2A1.1 enrichment correlated with H3K4me3 and H3K27ac, as well as with BRD4, H3.3 and Pol II binding (**Fig 2A, 2B & S5A, S5B Fig**), suggesting a role for mH2A1.1 in transcription initiation-regulated processes. At promoters, mH2A1.1 distribution inversely coincided with heterochromatin marks (**Fig 2A & S5A, S5B Fig**). We further determined that enrichment of mH2A1.1 centered at the TSS was proportional to the level of transcription (**Fig 2C-2D ; S5C-E Fig**). The profile of mH2A1.1 binding was greatest at the TSS of expressed genes and similar to the one of Pol II at the NFR, bordered by H3K27ac marked nucleosome regions, but larger than that of Pol II (**Fig 2E, S5F, S5G Fig**).

**Fig 2.**
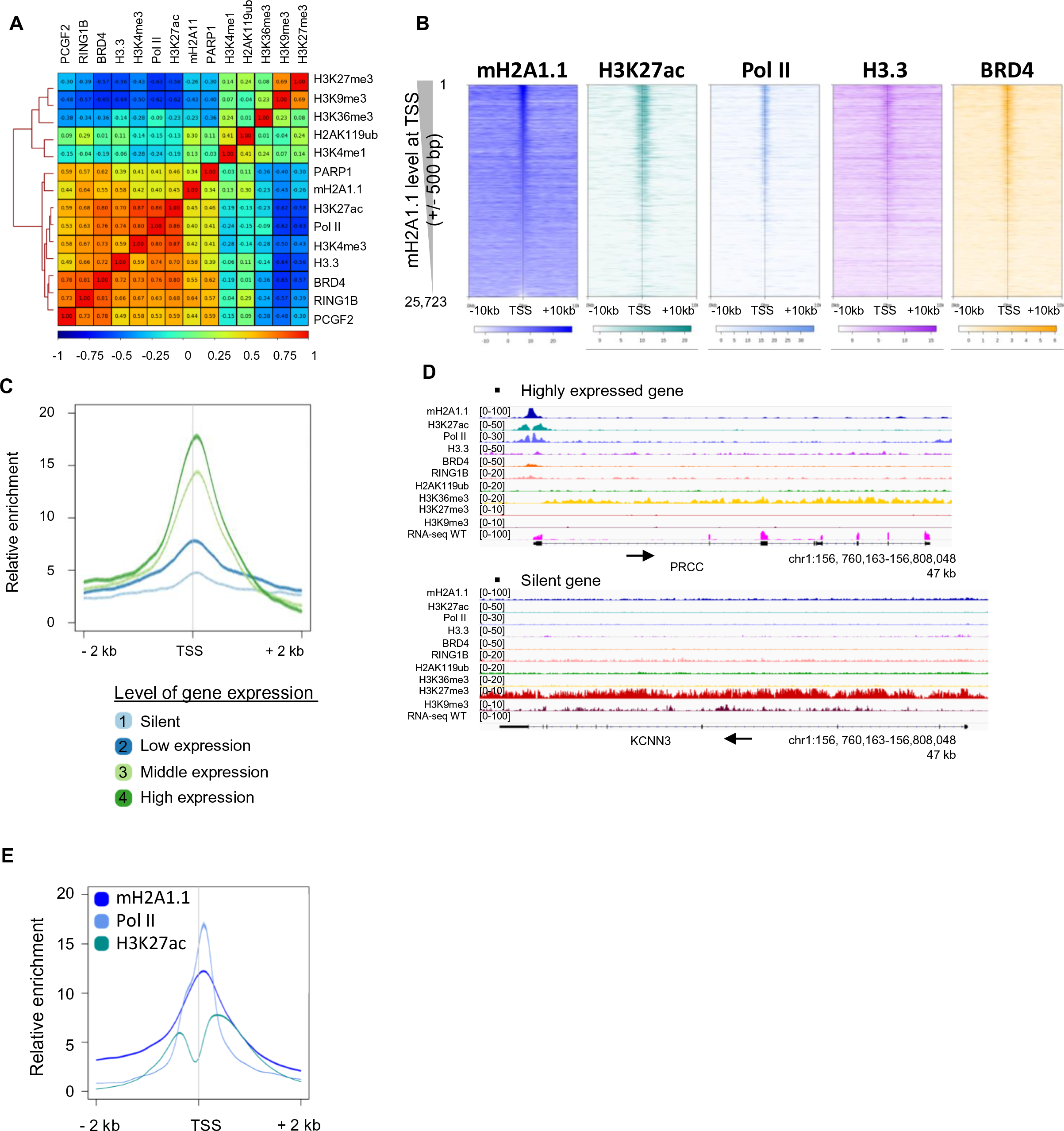
mH2A1.1 is recruited to promoters of active genes. (A) TSS (+/- 1kb) centered spearman correlation heatmap of ChIP-seq data. Correlations shown as in Fig1D. (B) Profiles of relative enrichment around the TSS (+/- 10 kb) of indicated ChIP-seq at human annotated genes (n= 25,723) ranked according to the mH2A1.1 level at TSS (+/-500bp). Colour intensity reflects level of ChIP-seq enrichment. Heatmaps are oriented with gene bodies placed on the right of the heatmap. (C) Metagene profiles of the average (+/-standard error) of mH2A1.1 enrichment at TSSs (+/- 2 kb) categorized in 4 groups according to gene expression levels measured using RNA-seq data. (D) Genome browser views of indicated ChIP-seq illustrating the binding of mH2A1.1 to the promoter region of a transcribed gene in an open chromatin state (top) and its absence on a silent gene in a closed chromatin state (bottom). Unstranded RNA-seq signal is also shown. Black arrows indicate direction of transcription. (E) Metagene profile of average (+/-standard error) of mH2A1.1, Pol II and H3K27ac enrichment at TSS (+/- 2kb) of transcribed genes (see Fig 2C, groups 2 to 4).

**Fig 3.**
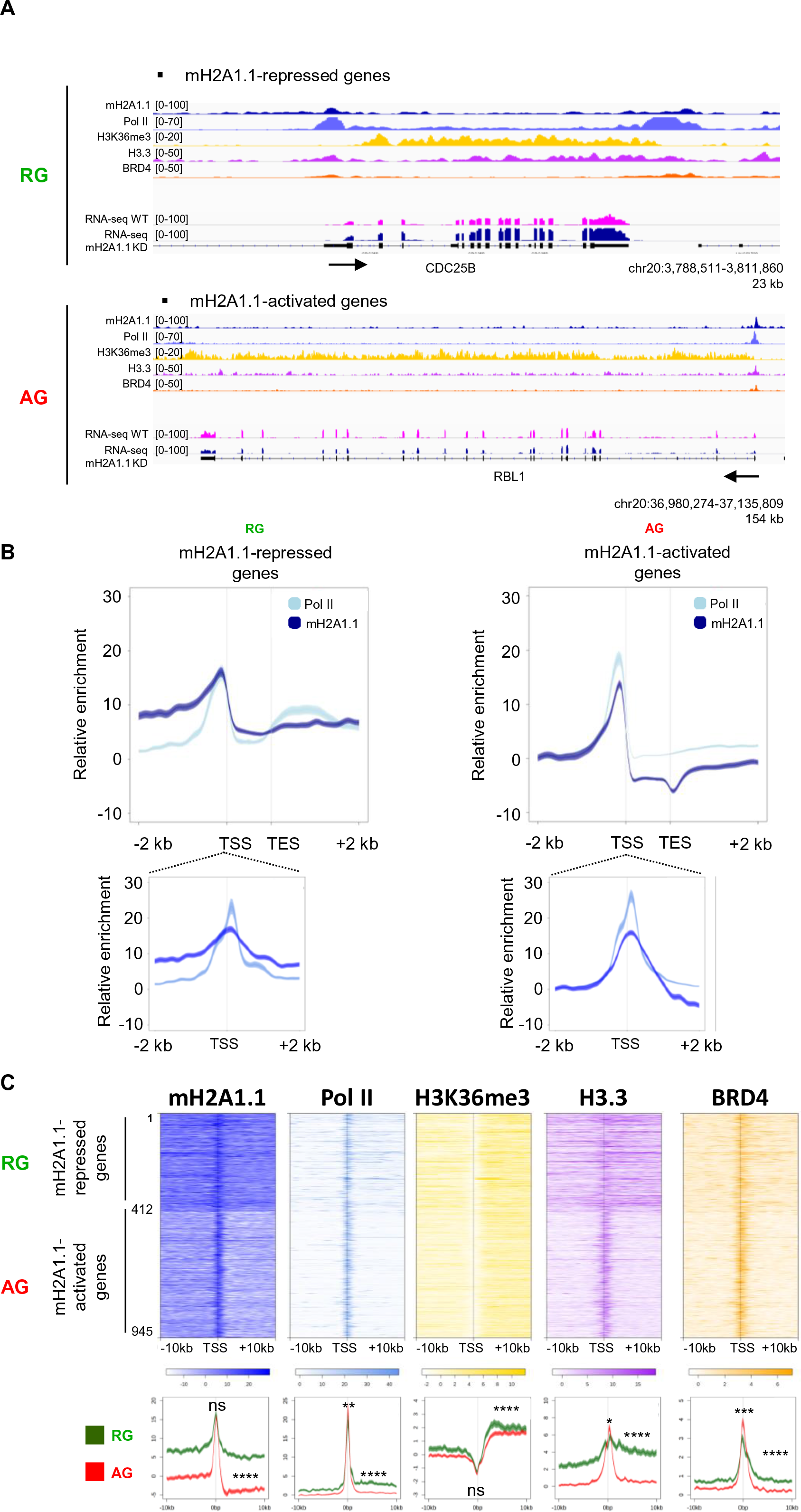
The chromatin landscapes of mH2A1.1-regulated genes. (A) Top panel: Heatmap profiles showing relative enrichment of indicated proteins and histone modifications around the TSS (+/- 10 kb) of mH2A1.1-regulated genes (see Fig 1A). On the top, mH2A1.1-repressed genes (1 to 412, n=412), on the bottom, mH2A1.1-activated genes (412 to 945, n=533). Color intensity reflects level of ChIP-seq enrichment. Heatmaps are oriented. Bottom panel: Metagene profiles of average (+/-standard error) of indicated ChIP-seq data around the TSS (+/- 10 kb) of mH2A1.1-regulated genes. Average profiles around the TSS of mH2A1.1-repressed genes are shown in green whereas average profiles around the TSS of mH2A1.1-activated genes are shown in red. Results of statistical difference analysis between these two groups are shown, either on the TSS (+/-50 bp) or on the gene body (+50 bp – TES). Complete statistical analysis is shown in S6 fig. ns, not significant. ** = p-value < 0.01, *** = p-value < 0.001, **** = p-value < 2.2x10^-16^. (B) Metagene profiles of average (+/-standard error) of mH2A1.1 and Pol II enrichment at mH2A1.1-regulated genes. (C) Genome browser views of ChIP-seq on a mH2A1.1-repressed gene (top) and a mH2A1.1-activated gene (bottom). Unstranded RNA-seq signals in control and mH2A1.1 KD are also shown. Black arrows indicate direction of transcription.

We next questioned whether this profile was linked to the mechanism by which mH2A1.1 regulates gene expression. To this end, we separately determined mH2A1.1 binding at genes repressed or activated (RG and AG) by mH2A1.1. At both gene categories, mH2A1.1 was highly enriched at the TSS (+/- 2 kb) (**Fig 3A-B**). However, mH2A1.1 binding was restricted to the TSS of mH2A1.1 AGs, while it associated both with promoter regions and the gene bodies of mH2A1.1 RGs **( Fig 3 )**. mH2A1.1 association correlated with the level of binding of Pol II, H3.3 and BRD4 **( Fig 3C ).** Interestingly at RGs, we detected Pol II at the promoter and over the elongation-characteristic H3K36me-marked gene body **( Fig 3B ; S6 Fig )**. At AGs, Pol II binding was essentially limited to the TSS. Thus, the Pol II distribution also discriminates both type of mH2A1.1-regulated genes. To note, although these genes are expressed, RING1B, PCGF2 and H2AK119ub, PRC1 subunits and associated modification, are present on RG-genomic regions (**S7 Fig**).

**Fig 4.**
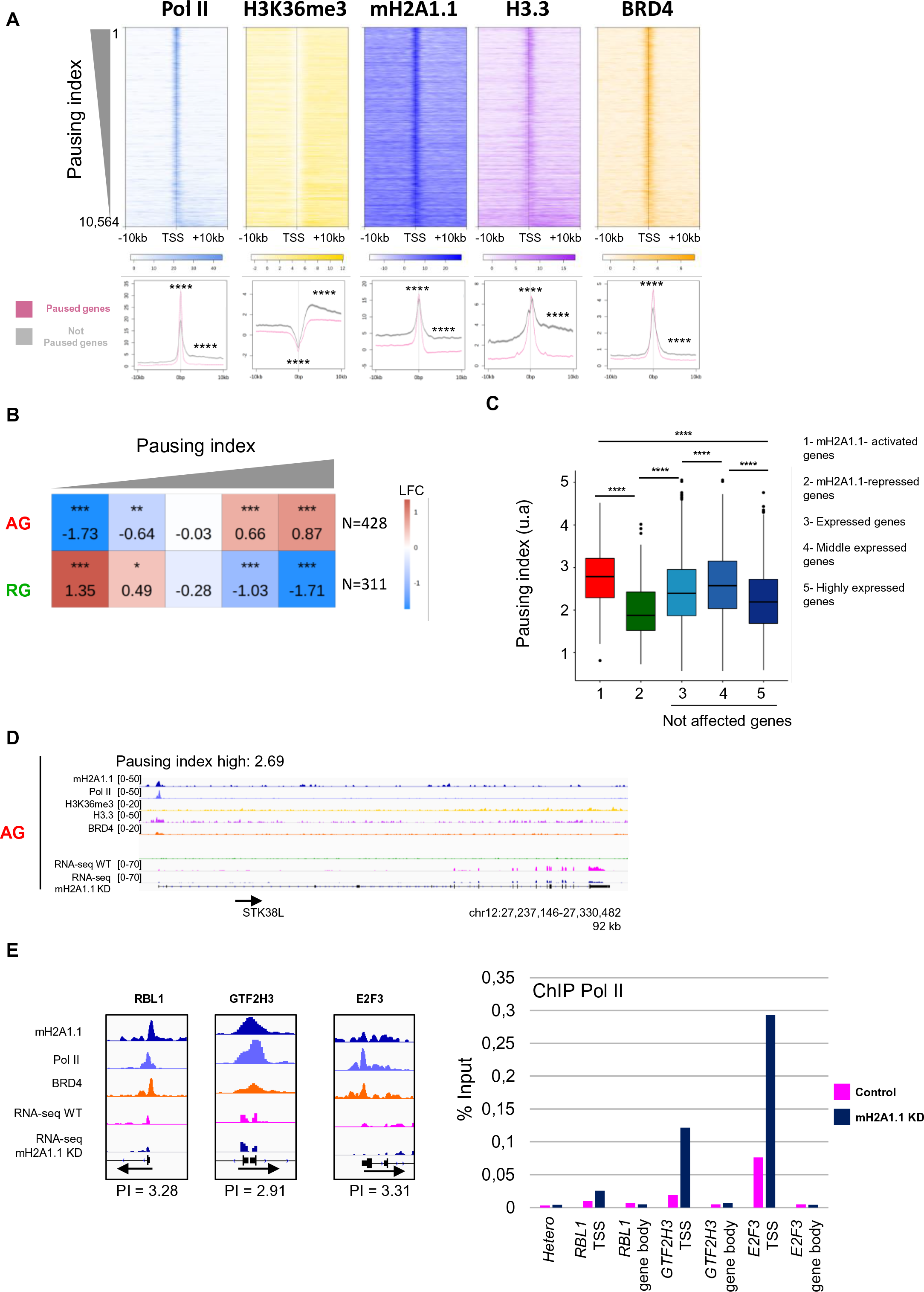
mH2A1.1-activated genes are regulated by Pol II pausing. (A) Top panel: Heatmap profiles showing enrichment of indicated factors and modifications around the TSS (+/- 10 kb) of transcribed genes (n=10,198) ranked by their pausing index. Colour intensity reflects level of ChIP-seq enrichment. Heatmaps are oriented. Bottom panel: Metagene profiles of average (+/-standard error) of indicated ChIP-seq data around the TSS (+/- 10 kb) of paused and not paused genes, as indicated in pink and black, respectively. Genes are considered as paused if their pausing index (PI) is > 2 (n=7,208). Genes are considered as a “not paused” if PI < 2 (n=3,356). Results of statistical difference analysis between these two groups are shown, either on the TSS (+/-50 bp) or on the gene body (+50 bp – TES). Complete statistical analysis is shown in S8A. **** = p-value < 2.2x10^-16^. (B) Fisher test heatmap showing enrichment of indicated mH2A1.1-target genes with genes divided in 5 equal size categories as a function of their pausing index. Stars indicate the significantly of the fisher exact tests; color map and values present in each scare highlight the log2 odd ratio (LOR) of the fisher exact test. N indicates the number of genes used for the analysis. (C) Boxplot comparing the pausing index of five indicated groups of genes. (Gene number in each group: 1, n=433, 2, n= 310, 3, n= 9.645, 4, n=5.176, 5, n=4.469). **** = p-value < 2.2x10^-16^. ns, not significant. Only genes characterized by a PI were used to generate this boxplot. (D) Genome browser view of indicated ChIP-seq on a paused gene. Unstranded RNA-seq signals in control and mH2A1.1 KD conditions are shown. Black arrows indicate direction of transcription. (E) Left panel: genome browser view of ChIP-seq of three mH2A1.1-activated genes (*RBL1, GTF2H3, E2F3*). Those genes are considered as paused genes with PI of 3,28, 2,9 and 3,2, respectively. Right panel: ChIP-qPCR of Pol II on WT and mH2A1.1-depleted cells. Hetero corresponds to a negative position. For each gene, Pol II enrichment was evaluated on the TSS and a gene body region. Results from additional biological replicates are given S10A.

**Fig 5.**
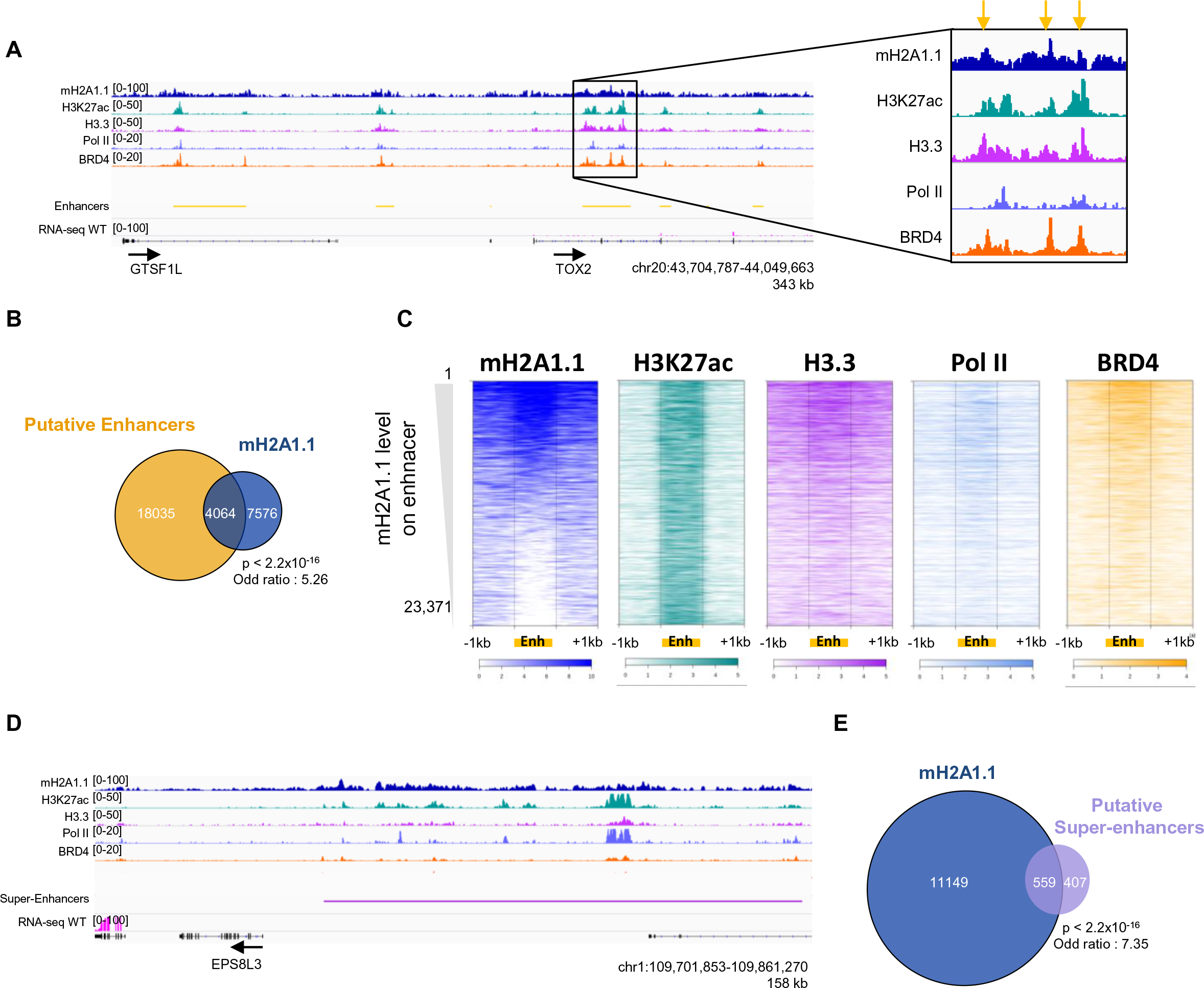
mH2A1.1 associates with enhancers and super-enhancers. (A) Genome browser view of indicated ChIP-seq illustrating occupancy of mH2A1.1 with “putative” enhancers. “Putative” enhancers are based on H3K27ac signal outside promoter regions using the ROSE package (Blinka et al., 2017). The black box shows a “zoom” on one enhancer. The orange arrows highlight the maximum signal of ChIP-seq data on this enhancer. (B) Overlap of “putative” enhancers with mH2A1.1 peaks. Enrichment of mH2A1.1-peaks with enhancers are measured using fisher exact tests. (C) Heatmap profiles showing indicated ChIP-seq data relative enrichment around the enhancers (+/- 1 kb). Colour intensity reflects the level of ChIP-seq enrichment. Each line represents an enhancer (from 1 to 23,371 enhancers). Enhancers are ranked according to the level of mH2A1.1 on enhancers, as indicated. (D) Genome browser view of indicated ChIP-seq illustrating occupancy of mH2A1.1 at “putative” super-enhancers (SEs). (E) Overlap of SEs with mH2A1.1 peaks. Enrichment of mH2A1.1-peaks with SEs are measured using fisher exact tests.

**Fig 6.**
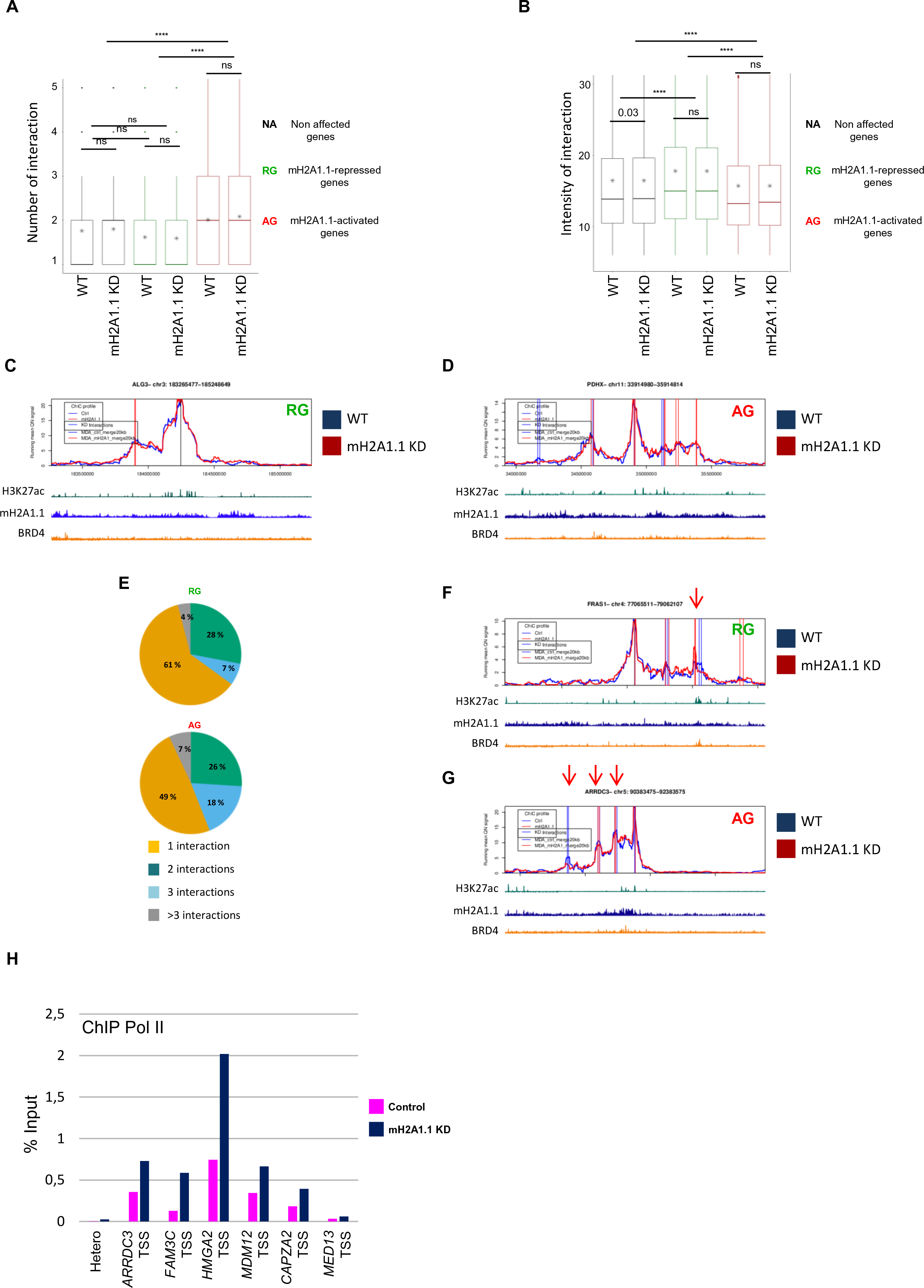
mH2A1.1 regulates gene expression within predefined 3D chromatin domains. (A) Boxplot showing the average number of PCHiC significant interactions per gene with adjacent genomic regions between genes not affected by mH2A1.1, mH2A1.1-repressed genes (n=181) and mH2A1.1-activated genes (n=282) in control and mH2A1.1 KD conditions. PCHiC significant interactions were determined using ChiCMaxima (Ben Zouari et al., 2019). Paired Wilcoxon test were used to compare control and mH2A1.1 KD conditions. ns, not significant. **** = p-value < 2.2x10^-16^. (B) Boxplot showing the mean of intensity of PCHiC interactions per gene between genes not affected by mH2A1.1, mH2A1.1-repressed genes (n=181) and mH2A1.1-activated genes (n=282) in control and mH2A1.1 KD conditions. (C) Snapshot of PCHiC data set on the mH2A1.1-repressed *ALO3* gene in control and mH2A1.1 KD conditions. Interaction intensity between the target gene and the associated genomic region are plotted over a 2 Mb gene domain around the promoter bait. Control (blue line) and mH2A1.1 KD (red line) are shown. The vertical bars correspond to PCHiC significant interactions conserved between two biological replicates in each condition. (D) Same as in (C) but for the mH2A1.1-activated *PDHX* gene. (E) Pie chart showing the percentage of mH2A1.1-target genes (RG on the top, AG on the bottom) having 1, 2, 3 or more than 3 PCHiC significant interactions. (F) Same as in (C) but this mH2A1.1-repressed gene, *FRAS1*, shows a reproducible gain of interaction with a specific genomic region (red arrow). (G) Same as in (D) but this mH2A1.1-activated gene, *ARRDC3*, shows a reproducible reduction of interaction with specific genomic regions (red arrow). (H) ChIP-qPCR of Pol II on WT and mH2A1.1-depleted cells. Hetero corresponds to a negative position. For each gene, Pol II enrichment was evaluated only on the TSS. Results from additional biological replicates are given S10B. On snapshots of PCHiC data, only results from replicate n°1 are shown here; second replicate n°2 in S14A-B.

**Fig 7.**
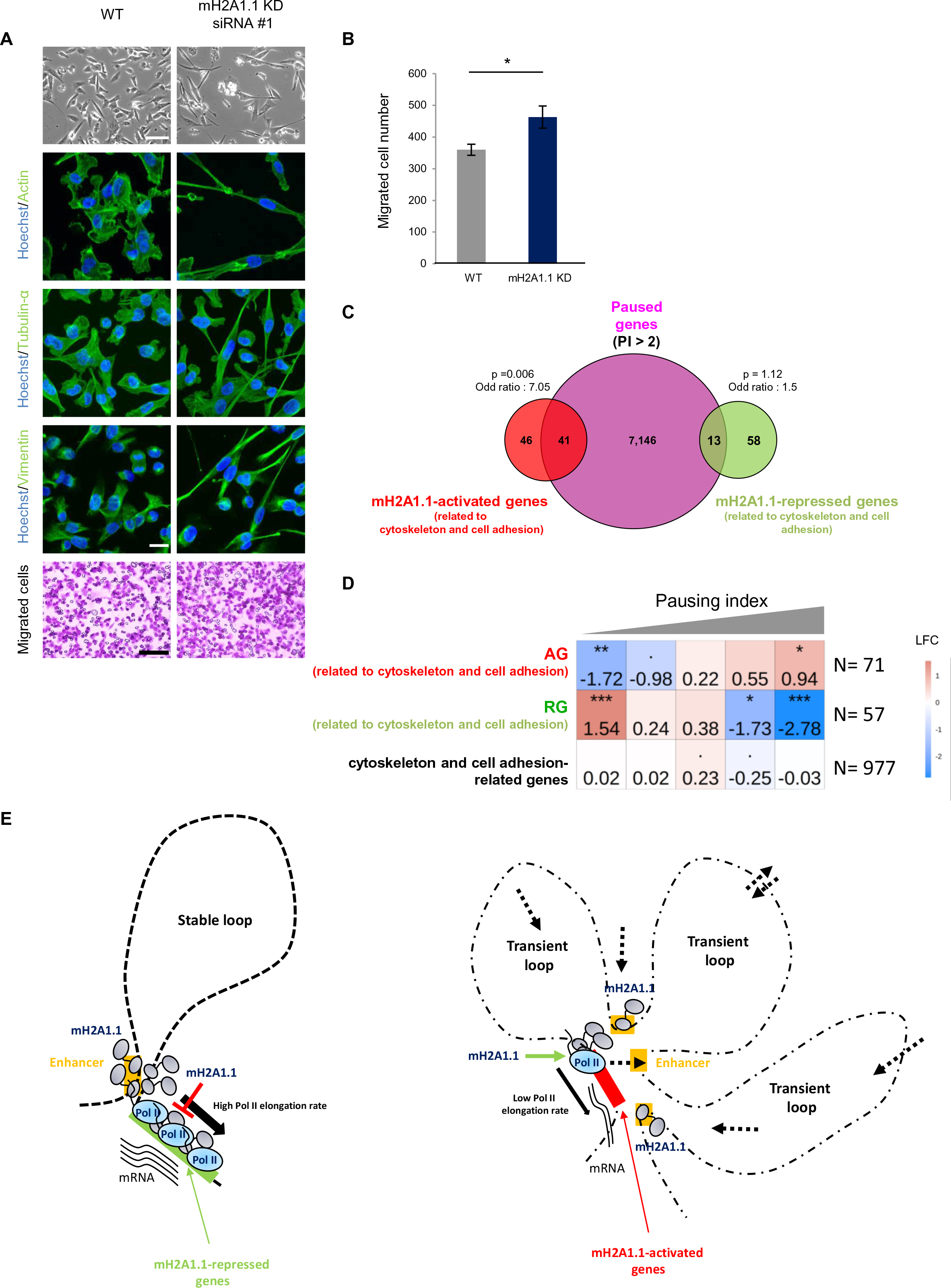
mH2A1.1 inhibits cell migration by favouring expression of paused genes involved in cytoskeleton and cell adhesion in MDA-MB231 cells. (A) Top: representative DIC microscopy images of WT and mH2A1.1 KD MDA-MB231 cells. Scale bar = 100 µM. Center: Immunofluorescence labelling of Actin, Tubulin-α and Vimentin. Nuclei are stained with Hoechst. Scale bar = 20 µM. Bottom: representative images of cells during Boyden chamber migration assay. Only migrated cells are labelled in purple. Scale bar = 150 µM. (B) Quantification of Boyden chamber assay presented in (A). Error bar represents s.d from n=3 independent experiments as illustrated in (A). “*” = p-value (p) < 0.05. (C) Overlap of paused genes (n=7,208) with mH2A1.1-regulated genes related to cytoskeleton and cell adhesion. Enrichment of this subgroup of mH2A1.1-regulated genes with paused genes are measured with fisher exact tests. (D) Fisher test heatmap showing enrichment of indicated mH2A1.1-target genes with genes divided in 5 equal size categories as a function of their pausing index. Stars indicate the significatively of the fisher exact tests; color map and values present in each scare highlight the log2 odd ratio (LOR) of the fisher exact test. N indicates the number of genes used for the analysis. (E) Working model describing the genomic organization of mH2A1.1-target genes. Left: mH2A1.1-repressed genes display a small number of stable interactions with enhancers or adjacent genomic regions characterized by bivalent chromatin marks. Pol II is enriched at the TSS and the gene body in this group of genes with a high Pol II elongation rate. The presence of mH2A1.1 all along the gene and associated enhancers slows Pol II elongation, maybe by favouring recruitment of repressors. Right: mH2A1.1-activated genes display a large number of transient interactions. Some of them are established with enhancers bound by BRD4 and possess a specific chromatin landscape. Pol II is mainly in pause in this group of genes, with a reduced Pol II elongation rate. Transient interactions between enhancers and promoters may promote Pol II pausing release, favoured by mH2A1.1.

### mH2A1.1 promotes gene expression of Pol II paused genes

The binding pattern of Pol II at mH2A1.1 AGs was reminiscent of Pol II paused genes (Adelman & Lis, 2012). To confirm this, we calculated the Pol II pausing index (PI) for transcribed genes using Pol II ChIP-seq data as described in (Adelman & Lis, 2012). Briefly, the pausing index corresponds to the ratio between total Pol II density in the promoter-proximal region (from -30 bp to + 300 bp around TSS) and total Pol II density in the transcribed region (from + 300 bp downstream TSS to TES). We plotted the ChIP-seq signal of Pol II and H3K36me3 around TSS +/- 10 kbp for each gene ranked according to their pausing index (**Fig 4A**). In agreement with the literature (Elrod et al., 2019), the level of H3K36me3 was greater over the body of genes with low PI compared to genes with high PI (**Fig 4A, and S8A**). We further observed that confinement of mH2A1.1 to the TSS and its absence from the gene body was characteristic of genes with a high PI (**Fig 4A, and S8A**). In agreement, the width of mH2A1.1 peaks overlapping with TSSs, as well as that of Pol II peaks, correlated negatively with the PI (**S8B-C Fig**). H3.3 follows the same binding profile as mH2A1.1. BRD4, RING1B and PARP1 were mainly enriched at the TSS, slightly more at high PI-genes (**Fig 4A, S8A and S9**).

The majority of mH2A1.1-AGs (85%) have a pausing index greater than 2 (**Fig S9C**), a PI value that can be used as a threshold to distinguish paused from not paused genes (Day et al., 2016). In agreement, the average PI of these genes was significantly higher than any other gene category tested (**Fig 4B-D**). At the difference, RGs are enriched in low PI-marked genes (**Fig 4B-C**). Furthermore, ChIPqPCR analysis of Pol II at three mH2A1.1-activated genes (*RBL1*, *GTF2H3* and *E2F3*) showed that the amount of Pol II at promoter regions was multiplied by a ∼3-fold factor upon siRNA reduction of mH2A1.1 (**Fig 4E and S10A**). On average, we observe that depletion of mH2A1.1 induces an increase in the PI of *RBL1*, *GTF2H3*, and *E2F3* genes by a factor of 1.4, 1.6, and 1.7, respectively. These results suggest that mH2A1.1 may enhance gene expression by promoting Pol II pause release.

### mH2A1.1 binds enhancers

In addition to promoters, mH2A1.1 also associates with intergenic regions (**Fig 1E**). Genome-wide, mH2A1.1 binding highly correlated with H3K4me1 (PCC of 0.55) and to a lesser extent with H3K27ac (PCC of 0.18), two chromatin marks which characterize enhancer regions (Creyghton et al., 2010) (**Fig 1D**). In agreement, we found that mH2A1.1 binding was significantly enriched at enhancers (Fisher exact test: p-value < 2.2x10^-16^ and odd ratio = 5.23) (**Fig 5A-B**). Enhancers bound by mH2A1.1 were further characterized by strong association of H3.3, Pol II, BRD4 and RING1B, which are marks of active enhancers (P. Chen et al., 2013; Lee et al., 2017) (**Fig 5C and S11A-B**). Strikingly, mH2A1.1-bound regions frequently formed large domains comprising a group of enhancers marked with H3K27ac (**Fig 5D**), which correspond to super-enhancers (Lovén et al., 2013; Whyte et al., 2013) (Fisher exact test: p-value < 2.2x10^-16^ and odd ratio = 7.35) (**Fig 5E and S11C**). Overall, these results show that mH2A1.1 binds enhancers and super-enhancers in association with BRD4 and RING1B.

### mH2A1.1-target genes regulation does not require changes in enhancer-promoter looping

Because mH2A1.1 binds enhancer and promoter regions (**Fig 3 and Fig 5**), an attractive hypothesis was that mH2A1.1 mediates chromatin folding. To test this hypothesis, we applied promoter capture HiC (PCHiC) using a collection of 19.225 promoter sequence fragments as bait (Schoenfelder et al., 2018) in WT and mH2A1.1 KD cells. Genomic interactions between promoters and other genomic regions were called by ChiCMaxima (Ben Zouari et al., 2019). We aggregated the total number of detected interactions per gene for mH2A1.1-activated, - repressed or -independent genes. For each category, the average number of interactions detected per gene was identical in control and mH2A1.1 KD cells (**Fig 6A**). The average intensity of these interactions with adjacent genomic regions (+/- 1.5 Mb around the gene) (**Fig 6B-D**) or of interactions with enhancer regions (**S12A Fig**) was also unaffected in the absence of mH2A1.1. Hence, chromatin interaction landscapes at mH2A1.1-regulated genes do not appear to require mH2A1.1. Yet, quantification of the PCHiC interactions showed that mH2A1.1-AGs have on average a greater number of interactions than mH2A1.1-RGs (**Fig 6A, 6E**). However, interactions at mH2A1.1-AGs showed weaker signal intensities (**Fig 6B, S12A**), suggesting that AG and RG reside within two types of interaction landscapes.

mH2A1.1 is enriched at the enhancers associated with RGs than those associated with AGs (**S12B-C**). Active chromatin marks and co-activators are also more abundant to RGs related enhancers than AGs related ones (H3.3, Pol II, BRD4) (**S12B-C, S13 Fig**) in agreement with the fact that these genes are in average more transcribed than AGs. Although, loss of mH2A1.1 did not induce any global changes in promoter contact numbers or frequencies, closer inspection of the interaction landscape of a few mH2A1.1-regulated genes revealed reproducible changes in the intensity of interactions at certain enhancers in mH2A1.1 KD vs wt cells (**Fig 6F, 6G** and **S14B-C**). For example, we observed an increase in the intensity of some interactions at the RG *FRAS1* upon mH2A1.1 depletion (**Fig 6F**), and a decrease in the intensity of interactions at the AG *ARDDC3* (**Fig 6G**). But this cannot be generalized, having also observed a decrease in the intensity of some interactions at RGs and an increase in the intensity of interactions at AGs (*data not shown*). Thus, the gain or loss of interactions does not appear to be related to transcriptional changes mediated by the loss of mH2A1.1. In addition, ChIP-qPCR of Pol II at the TSS of these AGs showed an increase in Pol II association at TSS upon loss of mH2A1.1 (**Fig 6H and S10B**). This observation is similar to that one observed for AGs (**Fig 4E**) whose 3D organization is not affected by mH2A1.1 depletion (**S14D Fig**). So, we conclude that chromatin looping does not interfere with the potential role of mH2A1.1 in Pol II release.

### mH2A1.1 inhibits cell migration by activating the expression of paused genes regulating cytoskeleton organization

mH2A1.1-target genes are involved in four main processes: cell cycle (9% of mH2A1.1-target genes), DNA repair (4%), cytoskeleton organization and cell adhesion (19%) (**S15A-B Fig and S4 Table**). The two first processes were expected based on earlier studies (Kim et al., 2018; Novikov et al., 2011; Judith C Sporn & Jung, 2012; Xu et al., 2012). However, the transcriptional role of mH2A1.1 on cytoskeleton organization and cell adhesion genes is poorly documented except the recent research of Marcus Buschbeck and colleagues showing that mH2A1.1 regulates those categories of genes in murine C2C12 cells (Hurtado-Bagès et al., 2020). Upon transfection of two different siRNA against mH2A1.1, MDA-MB231 cells became more elongated after 2-3 days (**Fig 7A and S16A**). Using immunofluorescence against cytoskeleton proteins (actin, tubulin-α, vimentin), we observed that the cytoskeleton organization was modified by the loss of mH2A1.1 (**Fig 7A**). Moreover, numerous mH2A1.1-regulated genes are involved in cell migration, such as *ARRDC3*, *SOCS4*, *HACE1* and *FBXL4* which are mH2A1.1-activated genes described as anti-migratory genes (Castillo-Lluva et al., 2013; Draheim et al., 2010; Mei et al., 2015; Stankiewicz et al., 2017) or *MMP14*, *EIF6*, *MT1E*, *JUND* and *DAPK3* which are mH2A1.1-repressed genes with pro-migratory properties (Cathcart et al., 2015; Kake et al., 2017; Pinzaglia et al., 2015; Ryu et al., 2012; Selvaraj et al., 2015). In agreement, upon depletion of mH2A1.1, the migratory capacity of MDA-MB231 cells was significantly increased compared to control cells (**Fig 7A-B**). The effect of knocking down the other isoform, mH2A1.2, was opposite to that of mH2A1.1 (**S16 Fig**).

Strikingly, mH2A1.1-AGs involved in cytoskeleton organization and cell adhesion were also amongst genes with a high Pol II pausing index (**Fig 7C, D**), compared to cell cycle and DNA repair mH2A1.1-AGs (**S15C Fig**). Overall, we conclude that mH2A1.1 impedes the migration capacity of MDA-MB231 breast cancer cells in part by promoting expression of genes modulating cell migration capacity.

## Discussion

Regulating gene expression in a particular cell type requires fine-tuning transcriptional response. The concentration and the relative ratio between factors required for these regulatory mechanisms ensure rapid adjustments to maintain homeostasis or to respond to stimuli and stress. In this study, we identify the histone variant mH2A1.1 as a means to operate these adjustments in TNBC cells.

We present the first genome-wide map of endogenous histone variant mH2A1.1 in human breast cancer cells. We discovered that the mH2A1.1 variant specifically associates with transcription regulatory elements, promoters and enhancers, in addition to large domains of facultative heterochromatin. Binding to promoters occurred in sharp, narrow peaks as opposed to the larger signals detected in heterochromatin seen previously (**Fig 2B and S4**)(H. Chen et al., 2014; Douet et al., 2017; Gamble et al., 2010; M. D. Lavigne et al., 2015). Moreover, we found that selective depletion of the mH2A1.1 isoform was sufficient to modify expression of hundreds of actively transcribed genes in the MDA-MB231 TNBC cell line (**Fig 1A-C**). All of these genes are highly expressed in this cell line. We uncovered two distinct mechanisms through which mH2A1.1 regulates their transcription and link them to the chromatin landscape in which the affected genes reside.

The first mechanism consists in dampening transcription of highly expressed genes. Indeed, in the absence of mH2A1.1, these mH2A1.1-RGs are overexpressed. mH2A1.1 binds the gene bodies alongside RNA pol II, as well as their associated promoters (**Fig 3 and S12B**). Importantly, these domains are also characterized by the presence of RING1B and the Polycomb-induced histone modification, H2AK119ub (Chan et al., 2018) on their enhancers and promoters (**Fig S7, S12, S13**). It is now recognized that the presence of Polycomb subunits does not always correlate with transcriptional repression states with the identification of specific PRC1 variants at a large number of active loci (Giner-Laguarda & Vidal, 2020) especially in cancer cells (Chan et al., 2018; Y. Zhang et al., 2020). Moreover, RING1B -target genes in MDA-MB231 cells are transcriptionally actives and highly expressed (Chan et al., 2018) as we found for mH2A1.1-targets genes (**Fig 1B,C**). We can postulate that the presence of mH2A1.1 may favor binding of the PRC1 complexes to moderate expression of mH2A1.1-RGs. Indeed expression levels at mH2A1.1-RGs are only slightly increased in mH2A1.1-depleted MDA-MB231 cells, similarly to observations in human lymphoma cell line (M. D. Lavigne et al., 2015), suggesting that mH2A1.1 may limit transcriptional noise and serve as a brake (**Fig 7E**).

The second mechanism is specific to genes in which RNA pol II is paused. Here, in contrast to the RGs, mH2A1.1 recruitment is restricted to the TSS of these genes. The deletion of mH2A1.1 leads to a reduction of the transcriptional level of these genes (mH2A1.1-AGs) as well as an accumulation of Pol II at their TSS (**Fig 1A-C, Fig 4E, Fig 6H**). It is indeed tempting to combine these two observations to propose that mH2A1.1 may assist in the conversion of promoter-locked RNA polymerase II into a productive and elongated Pol II. The presence of BRD4 at the TSS of mH2A1.1-AGs likely favors transcription elongation by playing a role in allosteric activation of P-TEFb (Winter et al., 2017). Then, we can imagine that mH2A1.1 could help to the recruitment of P-TEFb at promoter proximal region and thus contributes to the Pol II pause-release. Accumulation of Pol II at TSS could also be due to accumulation of torsional stress, through topoisomerase inhibition (Teves & Henikoff, 2014). BRD4 is known to overcome the torsional constraint to transcription by stimulating TOP1 activation concomitant with pause release events (Baranello et al., 2016). Deletion of mH2A1.1 could impair this process, resulting in the maintenance of torsional stress, accumulation of Pol II and inhibition of transcription. Finally, the chromatin organization at mH2A1.1-AGs, which are mostly characterized by Pol II pausing, seems more dynamic than at mH2A1.1-RGs, with more frequent but weaker contacts detected by PCHiC (**Fig 6**). In these domains, as Pol II is retained in pause at the TSS, its release could be facilitated by transient TSS-enhancer contacts in search for co-occupancy of coactivators and Pol II (**Fig 7E**). This observation can be generalized to all paused genes in the MDA-MB231 cell line (*data not shown*) identifying a new characteristic of paused genes with respect to their 3D organization of the genome. Conversely, the 3D organization of the mH2A1.1-RG loci appears to be relatively stable, reminiscent of a productive and well-organized environment for transcription (**Fig 6, Fig 7E**).

Currently, only one study has analyzed transcriptional activities of mH2A1.1 related to its genomic localization, and that is in murine muscle cell line C2C12 (Hurtado-Bagès et al., 2020). As in our study, although mH2A1.1 is mainly implicated in repression of transcription, a significant proportion of genes requires mH2A1.1 for their level of transcription. Among the activated genes, mH2A1.1 is enriched on genes encoding proteins related to adhesion, migration and the organization of extracellular matrix. But at the difference of our study, this recruitment occurs upstream of the TSS of these genes, the level of mH2A1.1 occupancy on the bodies of genes reduces with increasing transcriptional activity. However, Lavigne et al, identified a restricted recruitment of the mH2A1.2 isoform to TSS of mH2A1-regulated genes in two cancer cell types, HeLa and Namalwa cells (M. D. Lavigne et al., 2015) even if a comparison of genomic sites bound by mH2A1.2 nucleosomes revealed only a small overlap between HeLa and Namalwa cells. Similarly, restricted recruitments at TSS were also observed for mH2A1 and mH2A2 in human et mouse embryonic stem cells (Pliatska et al., 2018; Yildirim et al., 2014). Therefore, we believe that the mH2A variants are differentially recruited to regulatory sites depending on carcinogenic and differentiation state of the cells. In breast cancers, recruitment and thus the roles of mH2A1 variants must be subtype specific. The newly identified recruitment of mH2A1.1 would thus be TNBC specific, as not identified so far in luminal breast cancer cell lines (Gamble et al., 2010). It could then explain why we found a correlation between mH2A1.1 expression levels and survival rates only in TNBC patients (A.- C. Lavigne et al., 2014).

Interestingly, it is mainly genes involved in cell migration that are concomitantly regulated by the stimulatory activity of mH2A1.1 and Pol II released events (**Fig 7C-D**). Among mH2A1.1 AGs were anti-migratory genes such as ARRDC3 (Draheim et al., 2010), SOCS4 (Mei et al., 2015), HACE1 (Castillo-Lluva et al., 2013) and FBXL4 (Stankiewicz et al., 2017). A role for mH2A1.1 in the control of cell migratory capacities has already been identified in gastric cancer cells (F. Li et al., 2016), MDA-MB231 cells (Dardenne et al., 2012), and mouse cell lines (Dardenne et al., 2012; Posavec Marjanović et al., 2017). Here, in addition to that silencing of mH2A1.1 enhances cell migration in the MDA-MB231 cell line (**Fig 7A-B**), we begin to identify the underlying molecular mechanism with direct transcriptional stimulation by mH2A1.1 of genes highly dependent on the Pol II pausing (**Fig 7C-D**).

We further demonstrate that mH2A1.1-bound chromatin co-localizes with the H3K9me3 histone mark (**Fig S4A**). A fraction of these sites is devoid of H3K27me3 and could correspond to the identified mH2A localization at constitutive heterochromatin (Douet et al., 2017). However, the vast majority of mH2A1.1-bound H3K9me3-decorated chromatin contained also tri-methylated H3K27 (**Fig S4**). This difference may be a feature of the MDA-MB231 cell line, a high migratory capacity cancer cell line in which H3K9me3 histone marks are distributed unusually (Segal et al., 2018; Yokoyama et al., 2013). Overall, heterochromatin-related processes in this cancerous cell appear modified compared to non-cancerous cells and could potentially result from or favor malignant cellular transformation (Segal et al., 2018; Yokoyama et al., 2013). Thus, it could be interesting to further investigate additional molecular mechanisms, enzymes and epigenetic machineries altered in this cancer type.

Despite the association of mH2A1.1 with heterochromatin, its phenotypic knockdown was not sufficient to reactivate silenced genes present in these domains (**Fig 1B, 1C and S4**). Different hypothesis could explain this result. The first hypothesis could be that mH2A1 isoforms (mH2A1.1 and mH2A1.2) have redundant actions at heterochromatin. Here, we specifically depleted mH2A1.1 without affecting the expression of mH2A1.2 **( Fig S1D**). The presence of mH2A1.2 could be sufficient to maintain gene silencing, although mH2A1.2-occupied silent genes were also not reactivated upon mH2A1.2 knock-down (Dell’Orso et al., 2016). However, even if mH2A1 binding was shown to overlap with H3K27me3-decorated chromatin in primary human cells, no enrichment of H3K27me3 at mH2A1-regulated genes (both isoforms) was observed (H. Chen et al., 2014). The second hypothesis could be that mH2A1.1, as well as mH2A1.2, may serve as a lock to conserve heterochromatin stability/organization but are not required for gene silencing. In agreement with this hypothesis, two studies demonstrated that mH2A are implicated in the condensation of heterochromatin regions such as Lamin-Associated-Domains and repeated DNA elements without drastically affecting their expression level (Douet et al., 2017; Fu et al., 2015). Further analyses are needed to better characterize the role of mH2A at heterochromatin regions.

In this study, we demonstrate that a key function of mH2A1.1 is to orchestrate the proper transcriptional output of genes depending on their environment. Yet, mH2A1.1 did not seem necessary for chromatin topologies. We cannot exclude that mH2A1.1 participates in short-range or transient structural changes that our approach is not sensitive enough to identify. However, others have reported that key transcription factors or cofactors do not alter 3D folding and, in particular, enhancer-promoter looping (Schoenfelder & Fraser, 2019). For example, in MDA-MB231 cells, the Fra1 activator binds to promoters and enhancers, but does not mediate looping (Bejjani et al., 2021). In SEM leukemia cell lines, BRD4 inhibition has only minor effects on enhancer-promoter interactions (Crump et al., 2021) despite a strong effect on key-oncogenic target gene expression. Thus, stabilization of enhancer-promoter loops is not always a prerequisite for transcriptional fine-tuning by transcriptional regulators. We could speculate that their roles are more functional by facilitating interactions between enhancer-associated factors as it was observed for BRD4 through the formation of phase-separated condensates (Sabari et al., 2018). It would be interesting to test whether mH2A1.1 participates to this process especially since we have identified a preferential association of mH2A1.1 with SEs (**Fig 5D-E**). SEs are known to play an important part in many diseases, including several cancers in which they drive expression of oncogenes (Donati et al., 2018; Lovén et al., 2013) and the expression of mH2A1.1 is altered in cancer cells compared to normal tissues (Cantariño et al., 2013; A.-C. Lavigne et al., 2014; Judith C Sporn & Jung, 2012). SE function may be compromised by variations in mH2A1.1 level leading to inability to fine-tune transcriptional output, in particular via the first mechanism described above (**Fig 7E**) needed to avoid excessive transcription of oncogenes.

**S1 Fig.**
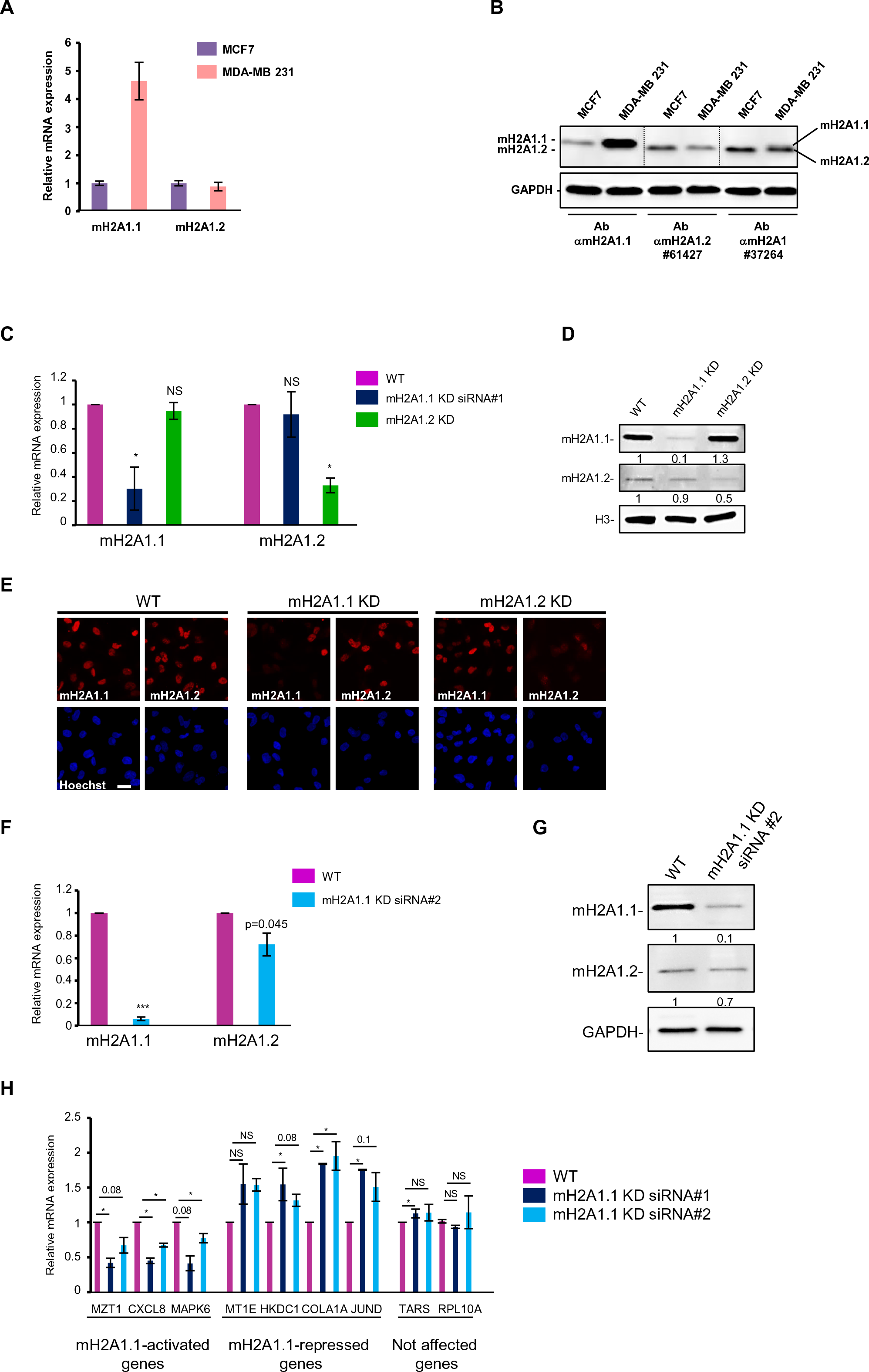
RNAi knock down of specific mH2A1 isoforms in MDA-MB231 cells. (A) RTqPCR on MDA-MB231 and MCF7 cells showing expression levels of mH2A1 isoforms. (B) Western blot on whole cell extracts of MDA-MB231 and MCF7 cells showing improved affinity of Ab αmH2A1.1 to recognize mH2A1.1 compared to Ab#37264 (Ab αmH2A1). Ab#61427 is specific to mH2A1.2. GAPDH is used as a loading control. (C) RTqPCR quantifying KD of mH2A1 isoforms. (D) Western blot showing specific depletion of mH2A1 isoforms protein. H3 is used as a loading control. (E) Immunofluorescence showing specific partial depletion of mH2A1 isoforms. DNA is labelled with Hoechst. Scale bar = 10 μm. (F) As in (C) but with a second siRNA against mH2A1.1 (siRNA #2). (G) As in (D) but with a second siRNA against mH2A1.1 (siRNA #2). GAPDH is used as a loading control. (H) RTqPCR analysis of a subset of RNAseq-defined mH2A1.1 regulated-genes. Genes are divided in three groups, as indicated. (C-H) Analysis were done three days post-transfection of specific siRNAs. RTqPCR, mRNA expressions are normalized by *RPLP0* mRNA. Error bars represent s.d from independent experiments (n>=2). “*” : p < 0.05, “***” : p < 0.001, ns, not significant. Quantifications are shown, normalized to protein loading control.

**S2 Fig.**
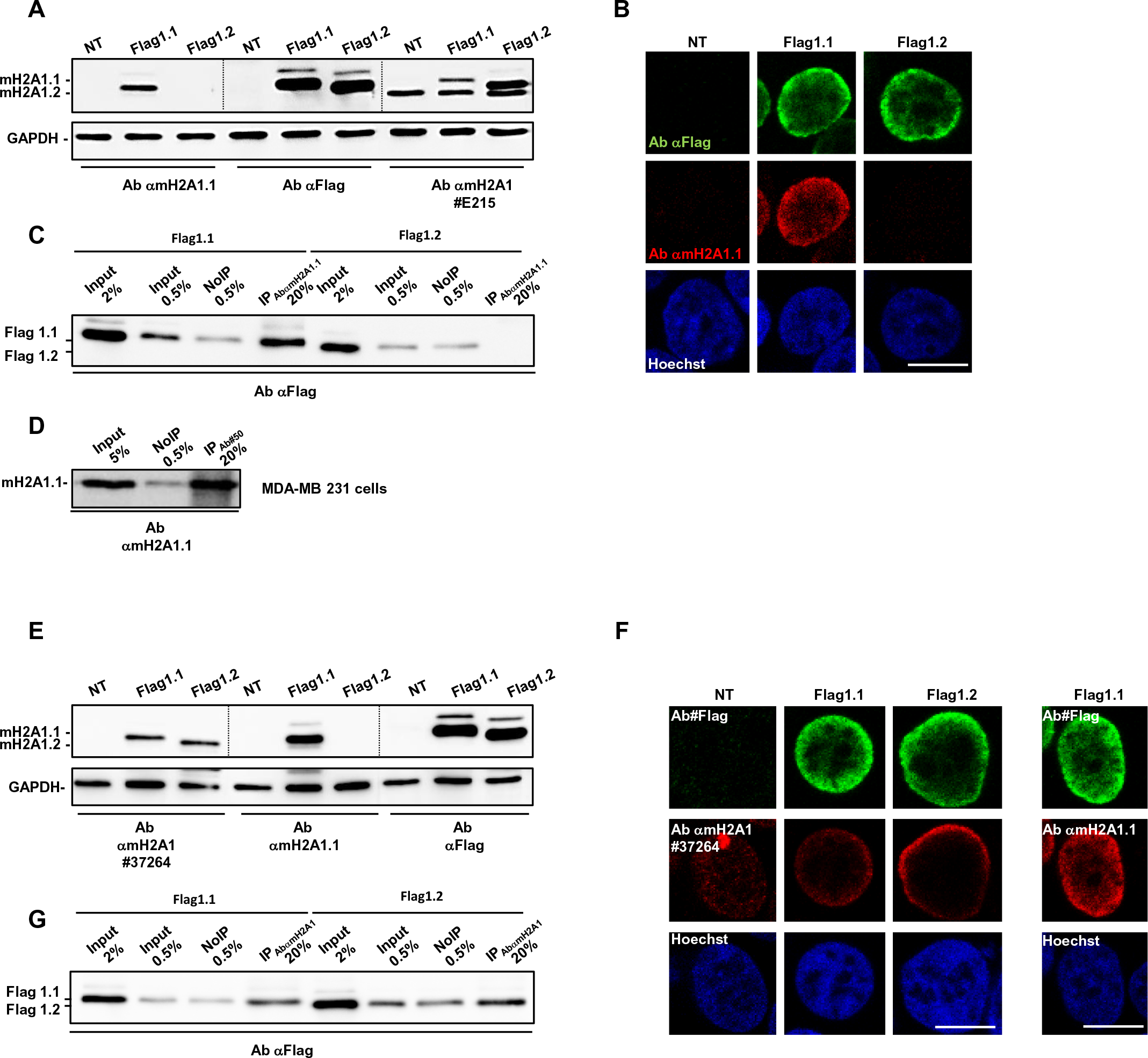
The antibody Ab αmH2A1.1 recognizes specifically the isoform mH2A1.1. (A) Western blot showing specific recognition of mH2A1.1 isoform by Ab αmH2A1.1 antibody. HEK-293T cells were transfected with plasmids coding for Flag-mH2A1.1 (Flag1.1) or Flag-mH2A1.2 (Flag1.2) fusion overexpressed-proteins. Western blot was then done with Ab αmH2A1.1, Ab#Flag and Ab#E215 (that preferentially recognizes mH2A1.2) antibodies on whole cell extracts. GAPDH is used as a loading control. (B) Immunofluorescence in HEK-293T cells showing specific recognition of mH2A1.1 isoform by Ab αmH2A1.1. (C) Western blot on ChIP extracts from HEK-293T cells overexpressing Flag1.1 or Flag1.2 showing that Ab αmH2A1.1 immunoprecipitates only mH2A1.1 isoform. Different extracts were loaded: Input fraction (Input), Non immunoprecipitated fraction (NoIP) and immunoprecipitated fraction (IP). Percentages represent fraction loaded on western blot compared to quantity used for ChIP. (D) Western blot showing that Ab αmH2A1.1 is also working in ChIP in MDA-MB231 cells on the endogenous protein. (E) As in (A), but for Ab αmH2A1 (Ab#37264) antibody showing that this antibody recognizes both isoforms but it less affine for Flag1.1 than Ab αmH2A1.1. (F) As in (B), but for Ab αmH2A1 (Ab#37264) antibody showing that this antibody recognizes both isoforms but it less affine for Flag1.1 than Ab αmH2A1.1. (G) As in (C) but for Ab αmH2A1 (Ab#37264) antibody showing that this antibody recognizes both isoforms but it less affine for Flag1.1 than Ab αmH2A1.1.

**S3 Fig.**
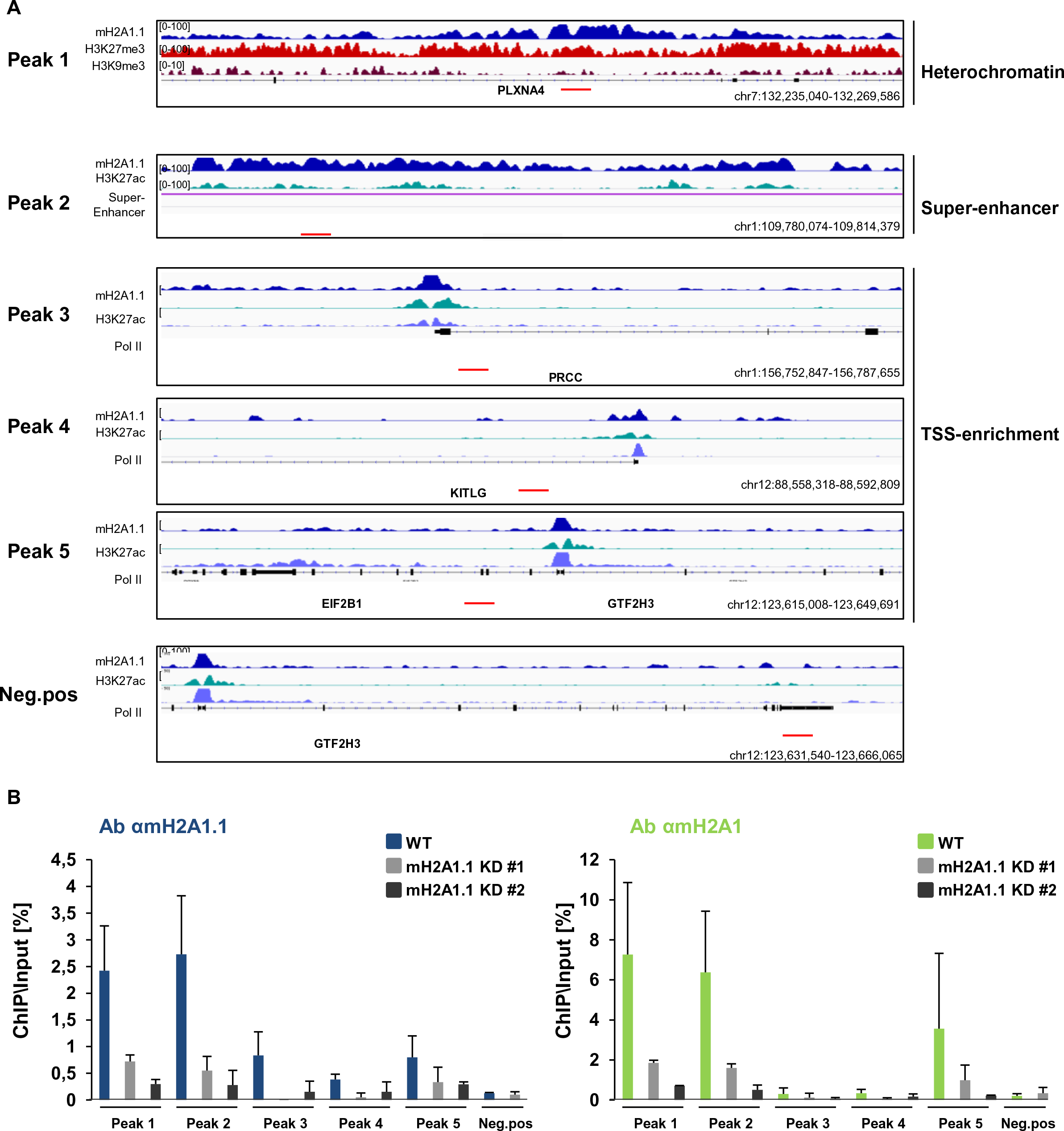
Validation of mH2A1 isoforms ChIP-seq by ChIP-qPCR on specific genomic loci. (A) mH2A1.1 binding at indicated genomic regions selected based on ChIP-sequencing. The black arrows show the direction of transcription. Genomic coordinates are also given. Localisation of primers used for ChIP-qPCR are shown in red. (B) Occupancy of mH2A1 isoforms (left part: Ab αmH2A1.1; right part: Ab αmH2A1) analysed by ChIP-qPCR in control cells (WT) and cells partially deficient for mH2A1.1 using two different siRNA (mH2A1.1 KD #1 and mH2A1.1 KD #2). Error bars represent +s.d from independent experiments (n>=2).

**S4 Fig.**
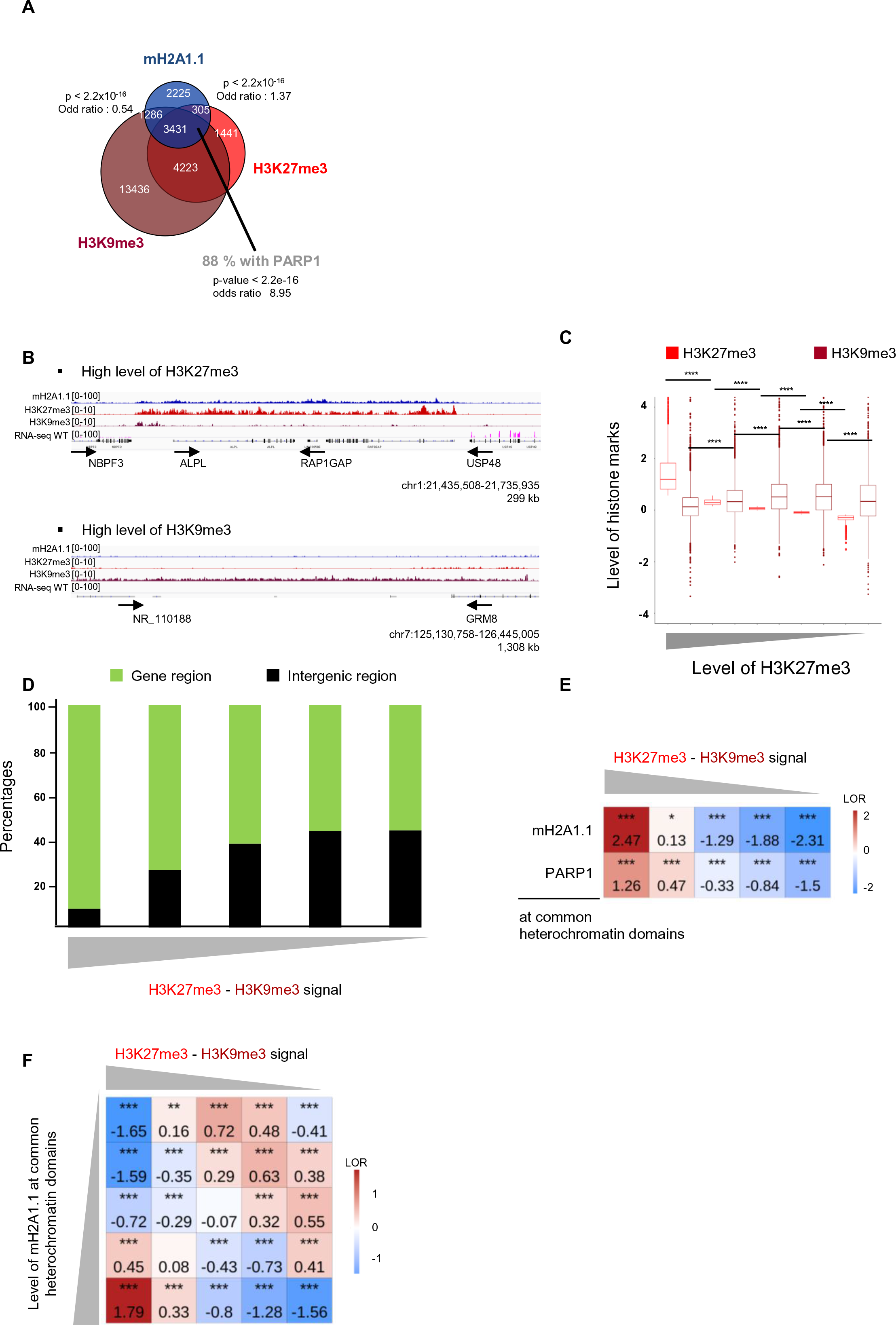
mH2A1.1 binds facultative heterochromatin domains. (A) Overlap of heterochromatin histone marks (H3K27me3 and H3K9me3) with mH2A1.1 peaks. Genome-wide enrichments of mH2A1.1 peaks with heterochromatin histone marks are measured with fisher exact tests. Enrichment of mH2A1.1 with PARP1 peaks was done on heterochromatin domains. (B) Genome browser view illustrating occupancy of mH2A1.1 with heterochromatin histone marks (H3K27me3 and H3K9me3). Top: region with high level of H3K27me3. Bottom: region with high level of H3K9me3. Unstranded RNA-seq signal is also shown. The black arrows show the direction of transcription. Genomic coordinates are given. (C) Boxplots showing H3K27me3 and H3K9me3 enrichment levels on H3K27me3-H3K9me3 common peaks. Common peaks were divided into 5 equal size categories according to the level of H3K27me3, as indicated. Statistical analyses to compare differences are given. “****” = p-value < 2.2x10^-16^. (D) Histogram showing proportions of heterochromatin (H3K27me3-H3K9me3 common peaks) on genomic regions (green) or intergenomic regions (black). Heterochromatin peaks were divided into 5 equal size categories according to difference between H3K27me3 and H3K9me3 signal, as mentioned (F) Fisher test heatmap showing enrichment of indicated ChIP-seq peaks (overlapping with common heterochromatin peaks) with heterochromatin peaks divided in 5 equal size categories as a function of differences between H3K27me3 and H3K9me3 signals. Stars indicate the significatively of the fisher exact tests; color map and values present in each scare highlight the log2 odd ratio (LOR) of the fisher exact test. (G) As for in (E) but this time mH2A1.1 peaks overlapping with heterochromatin domains are divided in 5 equal size categories as a function of mH2A1.1 signal.

**S5 Fig.**
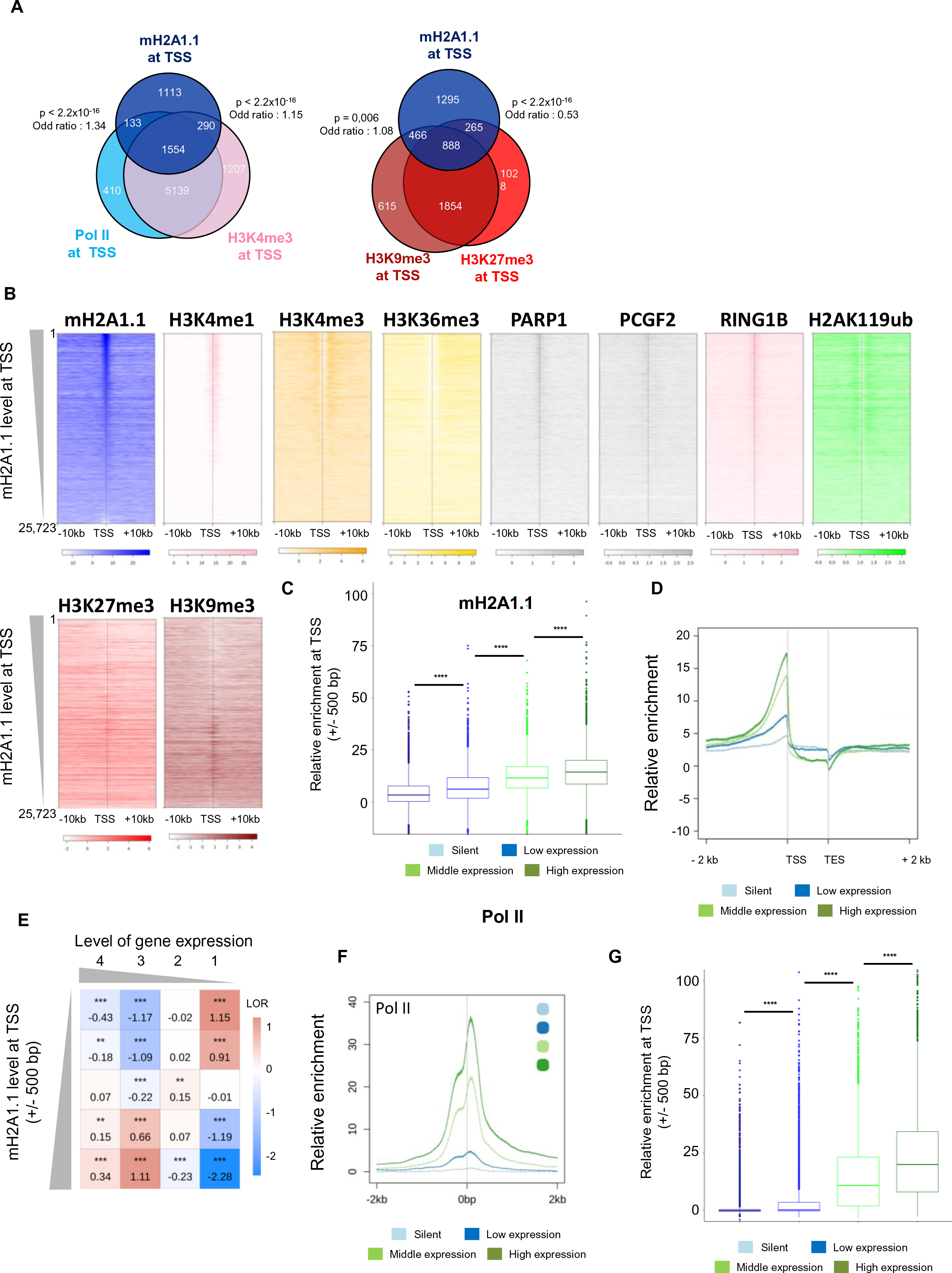
The mH2A1.1 isoform binds active promoters. (A) Overlaps of mH2A1.1 peaks with H3K4me3 and Pol II peaks at TSS (left) or H3K9me3 and H3K27me3 peaks (right). Enrichments of mH2A1.1 peaks with the ChIP-seq data at TSS are measured with fisher exact tests. (B) Heatmap profiles showing indicated ChIP-seq relative enrichment around the TSS (+/- 10 kb) at human annotated genes (n= 25,723) according to the level of mH2A1.1 at TSS (+/-500 bp). Colour intensity reflects level of ChIP-seq enrichment. Heatmaps are oriented. (C) Boxplots comparing the levels of mH2A1.1 at TSS (+/-500bp) of 4 groups of genes categorized by their gene expression levels, as indicated. “****” = p-value < 2.2x10^-16^. (D) Metagene plot of average (+/- standard error) of mH2A1 isoforms enrichment from TSS to TES (+/- 2 kb) categorized by gene expression levels, as indicated. (E) Fisher exact test heatmap showing the enrichment of mH2A1.1 at TSS (TSS +/-500 bp) on 4 groups of promoters categorized according to the level of gene expression. Legend as for Fig S4F. (F) as in (C) but for Pol II.

**S6 Fig.**
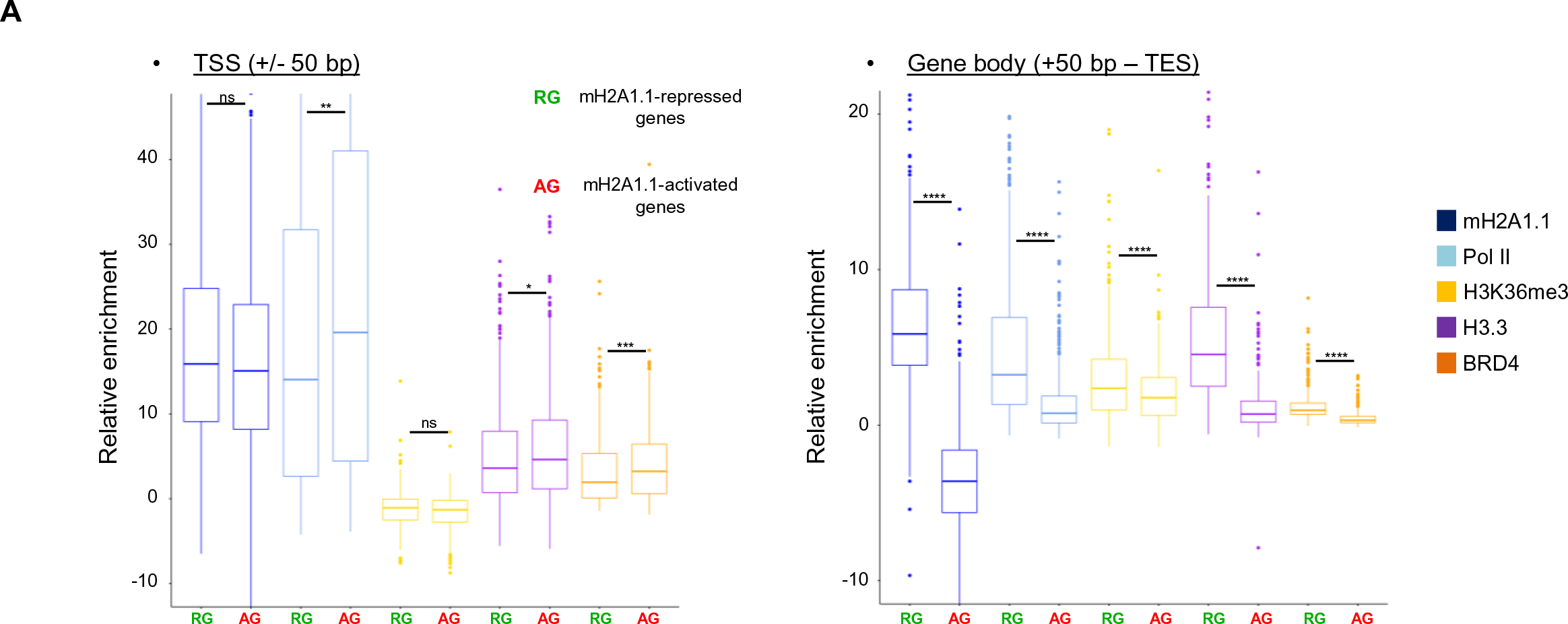
The mH2A1.1 isoform regulates expression of active genes in different chromatin environments. Boxplots showing the relative enrichment of indicated ChIP-seq between the mH2A1.1-repressed genes and mH2A1.1-activated genes. Left: enrichment on TSS (+/-50 bp); Right: relative enrichment of gene body (+50 bp to the TES). ns, not significant. “****” = p-value < 2.2x10^-16^.

**S7 Fig.**
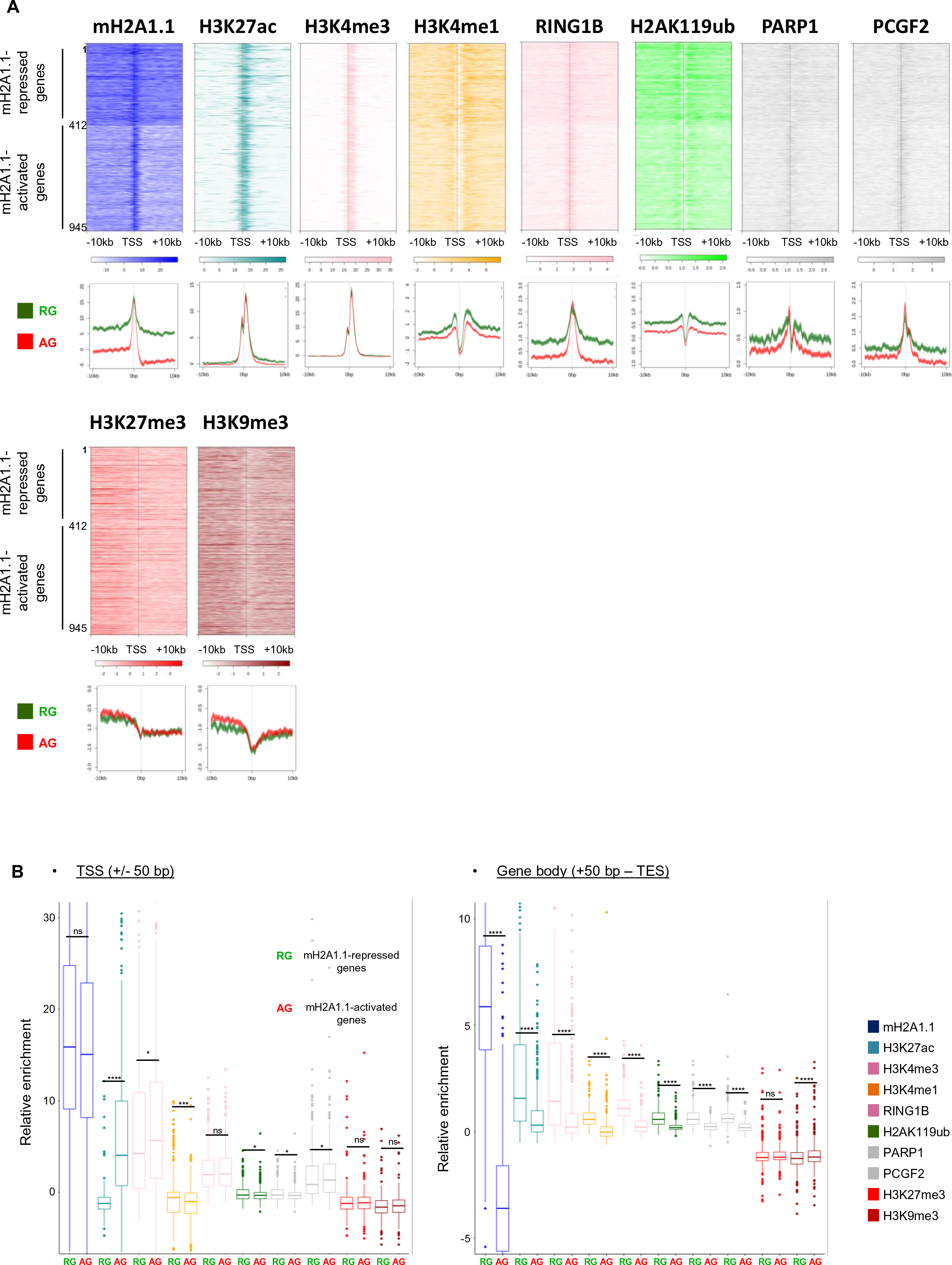
Chromatin environments of mH2A1.1-regulated genes. (A) Top panel: Heatmap profiles showing relative enrichment of indicated proteins and histone modifications around the TSS (+/- 10 kb) of mH2A1.1-regulated genes (see Fig 1A). On the top, mH2A1.1-repressed genes (1 to 412, n=412), on the bottom, mH2A1.1-activated genes (412 to 945, n=533). Color intensity reflects level of ChIP-seq enrichment. Heatmaps are oriented. Bottom panel: Metagene profiles of average (+/- standard error) of indicated ChIP-seq data around the TSS (+/- 10 kb) of mH2A1.1-regulated genes. Average profiles around the TSS of mH2A1.1- repressed genes are shown in green whereas average profiles around the TSS of mH2A1.1- activated genes are shown in red. (B) Boxplots showing the relative enrichment of indicated ChIP-seq between the mH2A1.1-repressed genes and mH2A1.1-activated genes. Left: enrichment on TSS (+/- 50 bp); Right: relative enrichment of gene body (+50 bp to the TES). ns, not significant. “****” = p-value < 2.2x10^-16^.

**S8 Fig.**
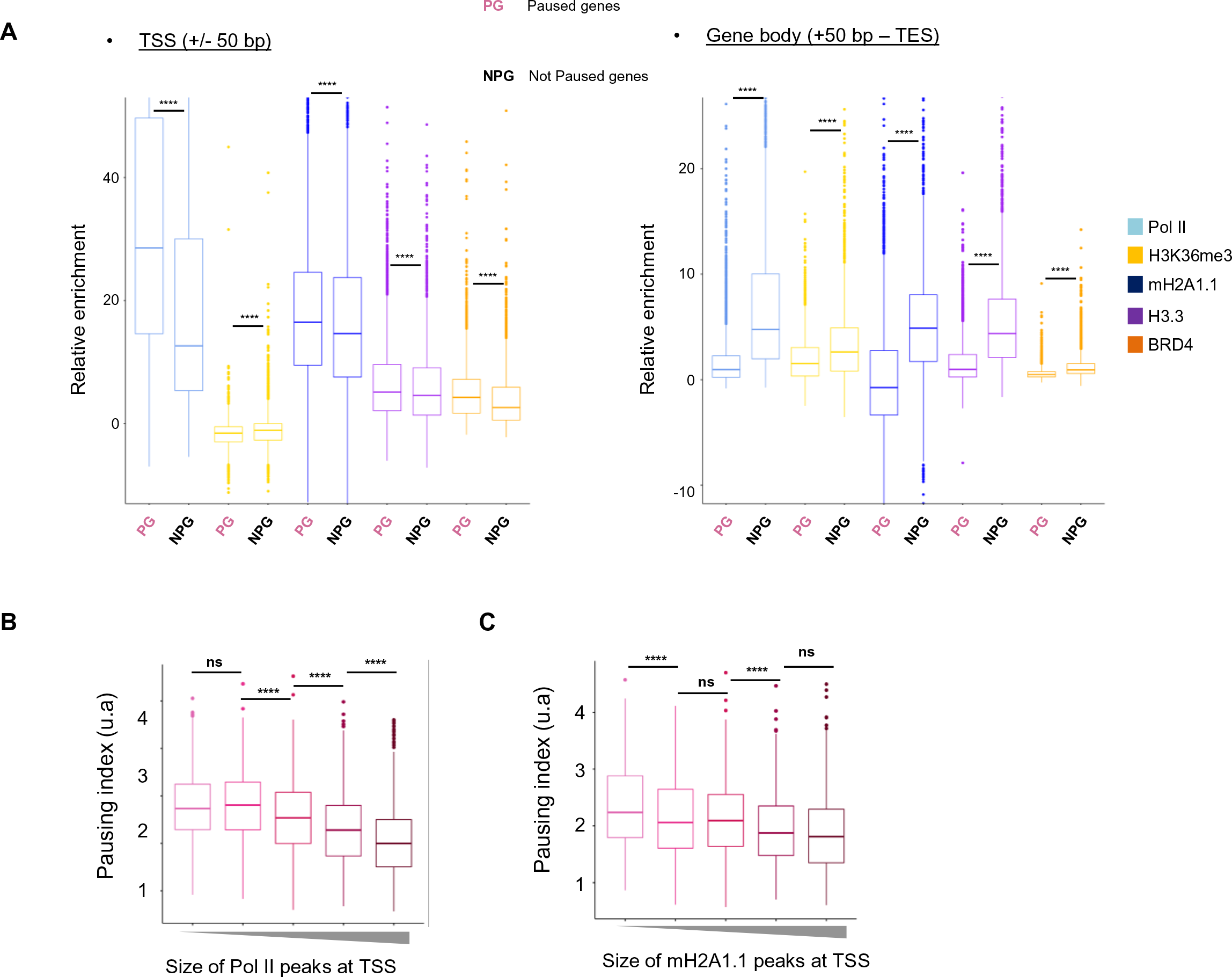
Chromatin environments of paused genes. (A) Boxplots showing the relative enrichment of indicated ChIP-seq between paused and not paused genes. Genes are considered as paused if their pausing index (PI) is > 2 (n=7,208). Genes are considered as a “not paused” if PI < 2 (n=3,356). Left: boxplot showing the relative enrichment of ChIP-seq around TSS (+/- 50 bp). Right: boxplot showing the relative enrichment of ChIP-seq on the gene body (from + 50bp to TES). ns, not significant. “****” = p-value < 2.2x10^-16^. (B) Boxplot comparing the pausing index of 5 categories of Pol II-bound genes divided according to the width of Pol II peaks. ns, not significant. “****” = p-value < 2.2x10^-16^. (C) Same as in (B) but for mH2A1.1-bound genes.

**S9 Fig.**
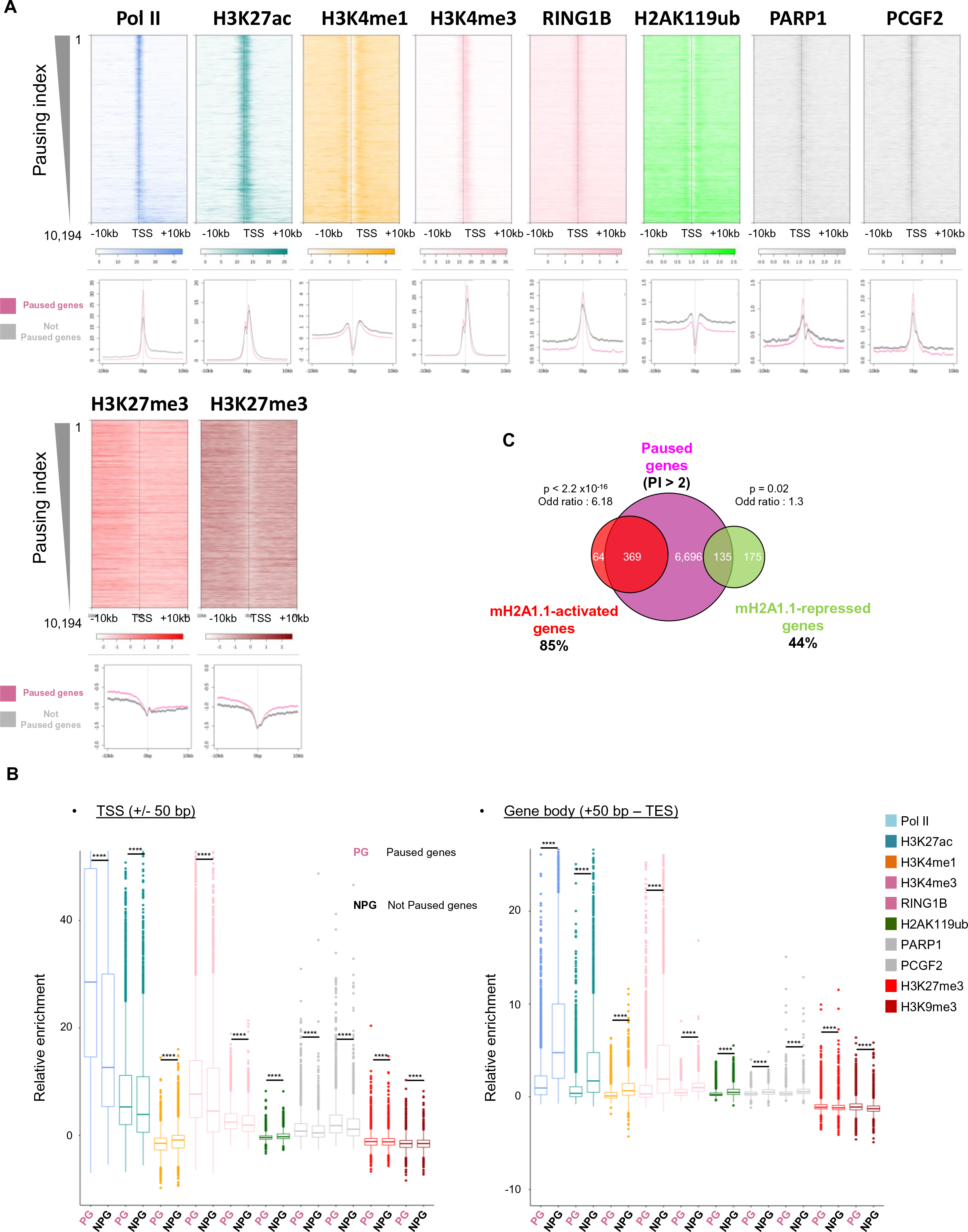
Chromatin environments of paused genes. (A) Top panel: Heatmap profiles showing enrichment of indicated factors and modifications around the TSS (+/- 10 kb) of transcribed genes (n=10,198) ranked by their pausing index. Colour intensity reflects level of ChIP-seq enrichment. Heatmaps are oriented. Bottom panel: Metagene profiles of average (+/- standard error) of indicated ChIP-seq data around the TSS (+/- 10 kb) of paused and not paused genes, as indicated in pink and black, respectively. Genes are considered as “paused” if their pausing index (PI) is > 2 (n=7,208). Genes are considered as a “not paused” if PI < 2 (n=3,356). (B) Boxplots showing the relative enrichment of indicated ChIP-seq between paused and not paused genes. Genes are considered as paused if their pausing index (PI) is > 2 (n=7,208). Right: boxplot showing the relative enrichment of ChIP-seq on the gene body (from + 50bp to TES). ns, not significant. “****” = p-value < 2.2x10^-16^. (C) Overlap of mH2A1.1- regulated genes with paused genes. Enrichment of mH2A1.1-target genes with paused genes are measured using fisher exact tests. Of note, only mH2A1.1-target genes characterized by a PI were used to generate this Venn diagram.

**S10 Fig.**
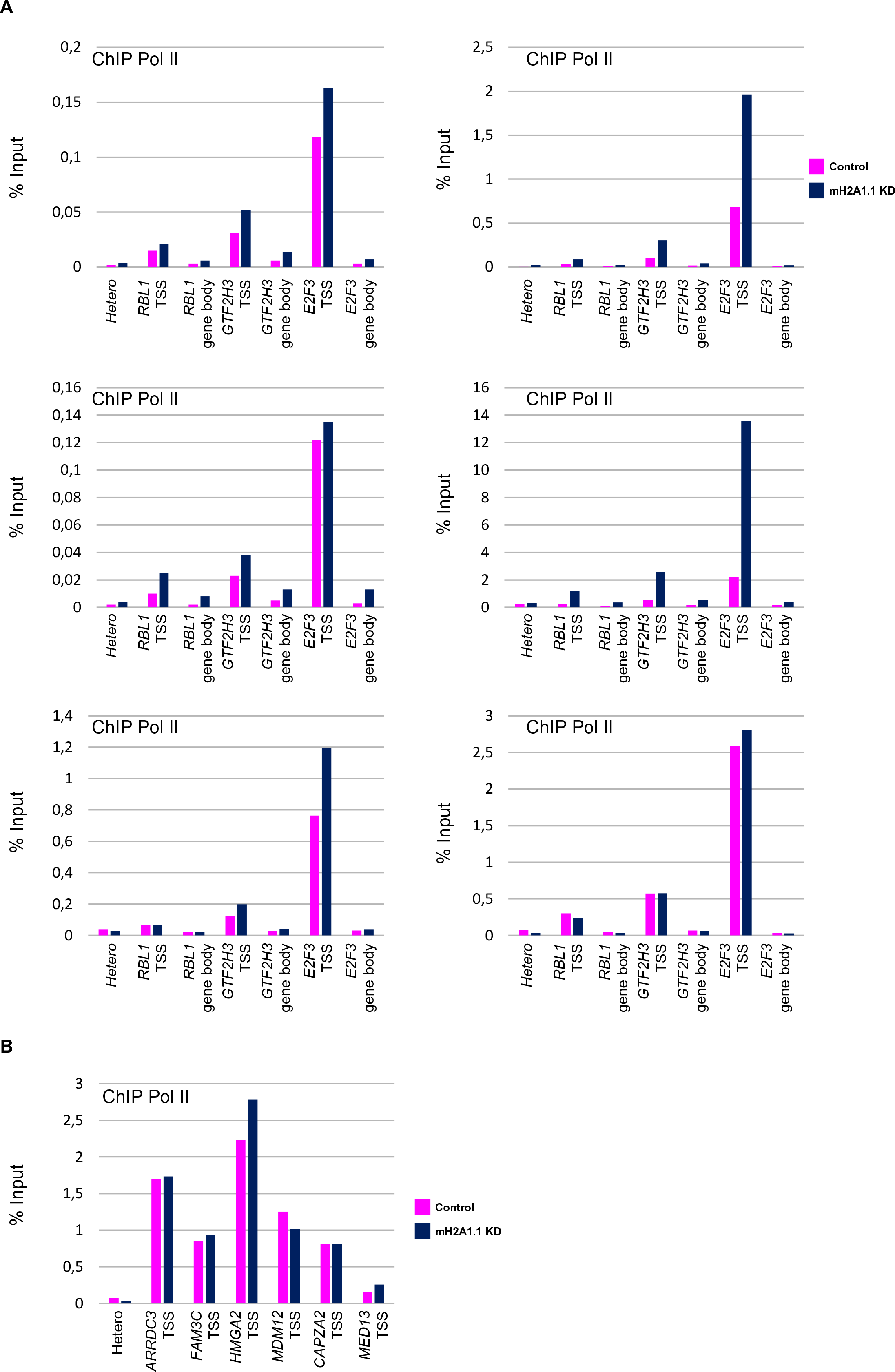
mH2A1.1 favours Pol II pausing release. (A) Biological replicates of ChIPqPCR of Pol II in control and mH2A1.1 KD conditions. The first biological replicate is shown Fig 4E. (B) ChIPqPCR of Pol II in control and mH2A1.1 KD conditions on mH2A1.1-activated genes that lose interactions with adjacent genomic regions. The first biological replicate in shown Fig 6H.

**S11 Fig.**
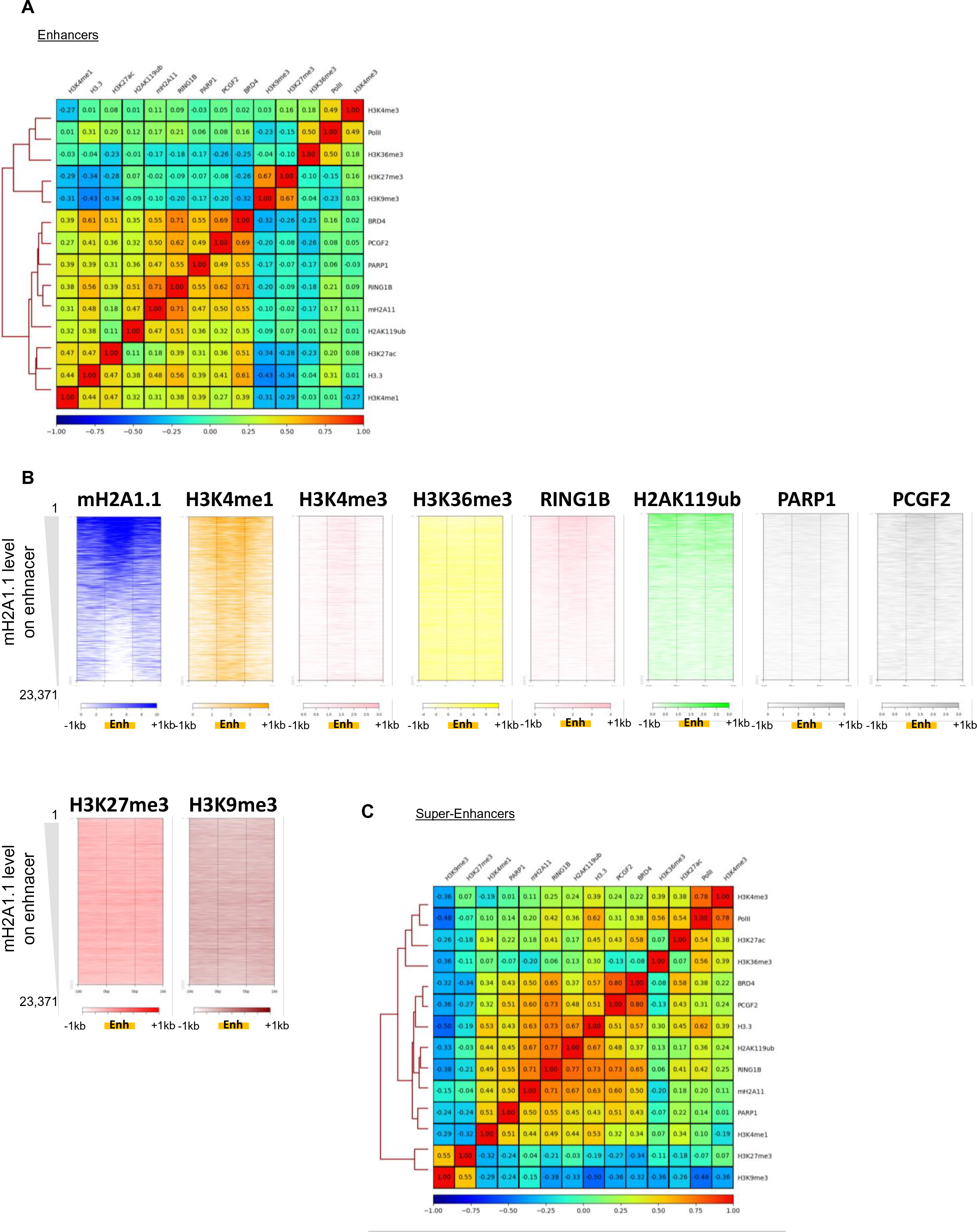
The mH2A1.1 isoform binds active enhancers and super-enhancers. (A) “Putative” enhancers centered spearman correlation heatmap of ChIP-seq data. Correlations shown as in Fig1D. Enhancers are based on H3K27ac signal outside promoter regions using the ROSE package (Blinka et al., 2017). (B) Heatmap profiles showing indicated ChIP-seq data relative enrichment around the enhancers (+/- 1 kb). Colour intensity reflects the level of ChIP-seq enrichment. Each line represents an enhancer (from 1 to 23,371 enhancers). Enhancers are ranked according to the level of mH2A1.1 on enhancers, as indicated. (C) Same as in (A) but on “putative” super-enhancers. SEs were defined using the ROSE package based on H3K27ac signal outside promoters regions (Blinka et al., 2017).

**S12 Fig.**
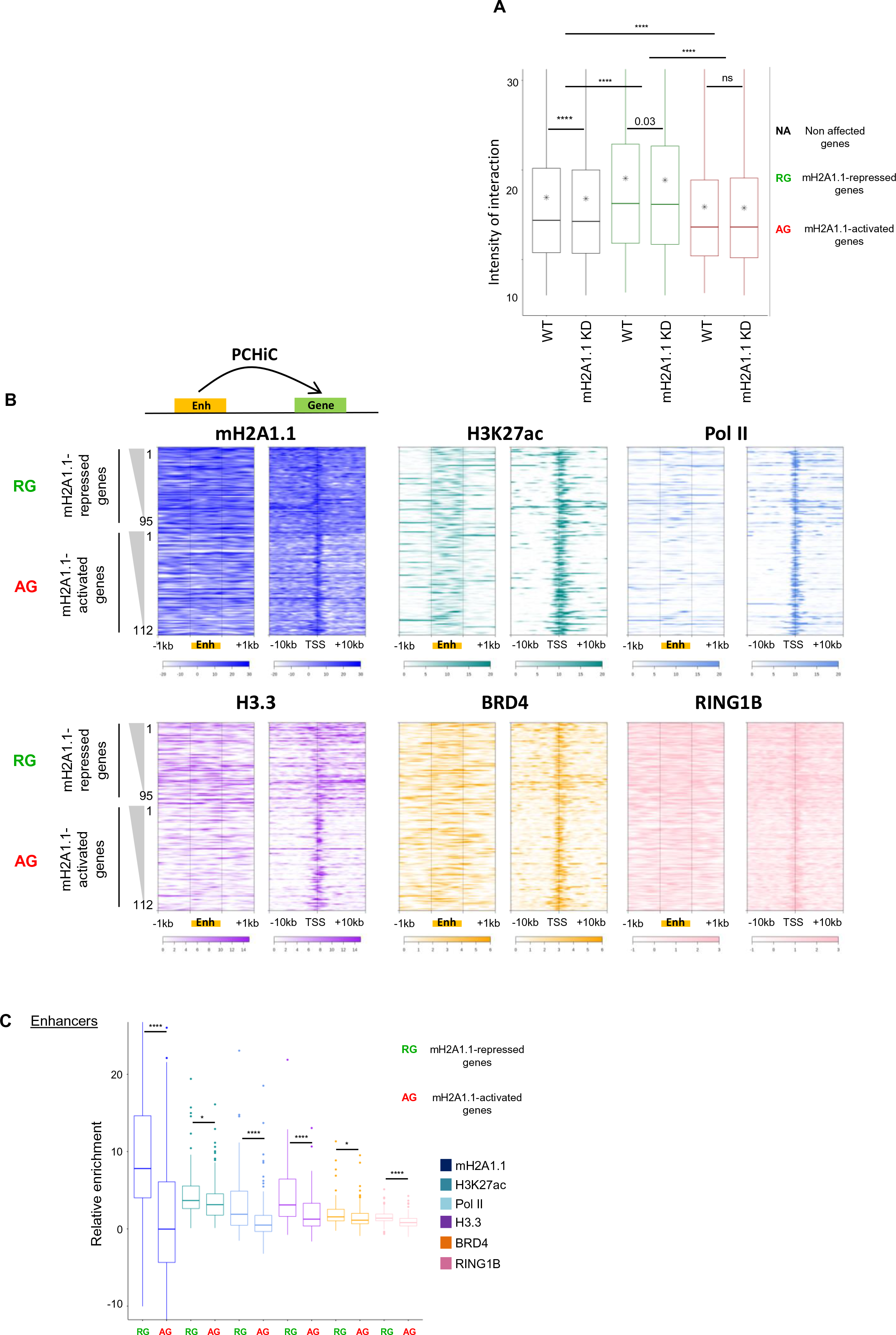
The chromatin environment at enhancers of mH2A1.1-regulated genes. (A) Boxplot showing the intensity of PCHiC interactions between mH2A1.1-non affected genes, mH2A1.1- repressed genes (n=181) and mH2A1.1-activated genes (n=282) in control and mH2A1.1 KD conditions with their respective enhancers. Enhancers of mH2A1.1-regulated genes were determined using PCHIC data and enhancers annotations (Materials and Methods). ns, not significant. “****” = p-value < 2.2x10^-16^. Paired Wilcoxon test were used to compare control and mH2A1.1 KD conditions. (B) Heatmap profiles showing indicated ChIP-seq data relative enrichment around the TSS (+/- 10 kb) of mH2A1.1-regulated genes (right) and their associated enhancers (+/- 1kb) (left). Enhancers of mH2A1.1-regulated genes were determined using PCHIC data and enhancers annotations (Materials and Methods). More than one enhancer can significantly be in interaction with mH2A1.1-regulated genes. To simplify, only one enhancer per gene was randomly conserved to generate those heatmaps. Top: mH2A1.1-repressed genes (1 to 95). Bottom: mH2A1.1-activated genes (1 to 112). Genes are ranked according to their expression level differences between the control and mH2A1.1 KD conditions. Some mH2A1.1-target genes are not present in those heatmaps because they do not have PCHiC significate interactions with an enhancer or are not present in the PCHiC database. Colour intensity reflects level of ChIP-seq enrichment. TSS-centered heatmap profiles are orientated. (B) Boxplots comparing the relative enrichment of indicated ChIP-seq between the enhancers of mH2A1.1-repressed genes and the enhancers of mH2A1.1- activated genes. ns, not significant. “****” = p-value < 2.2x10^-16^.

**S13 Fig.**
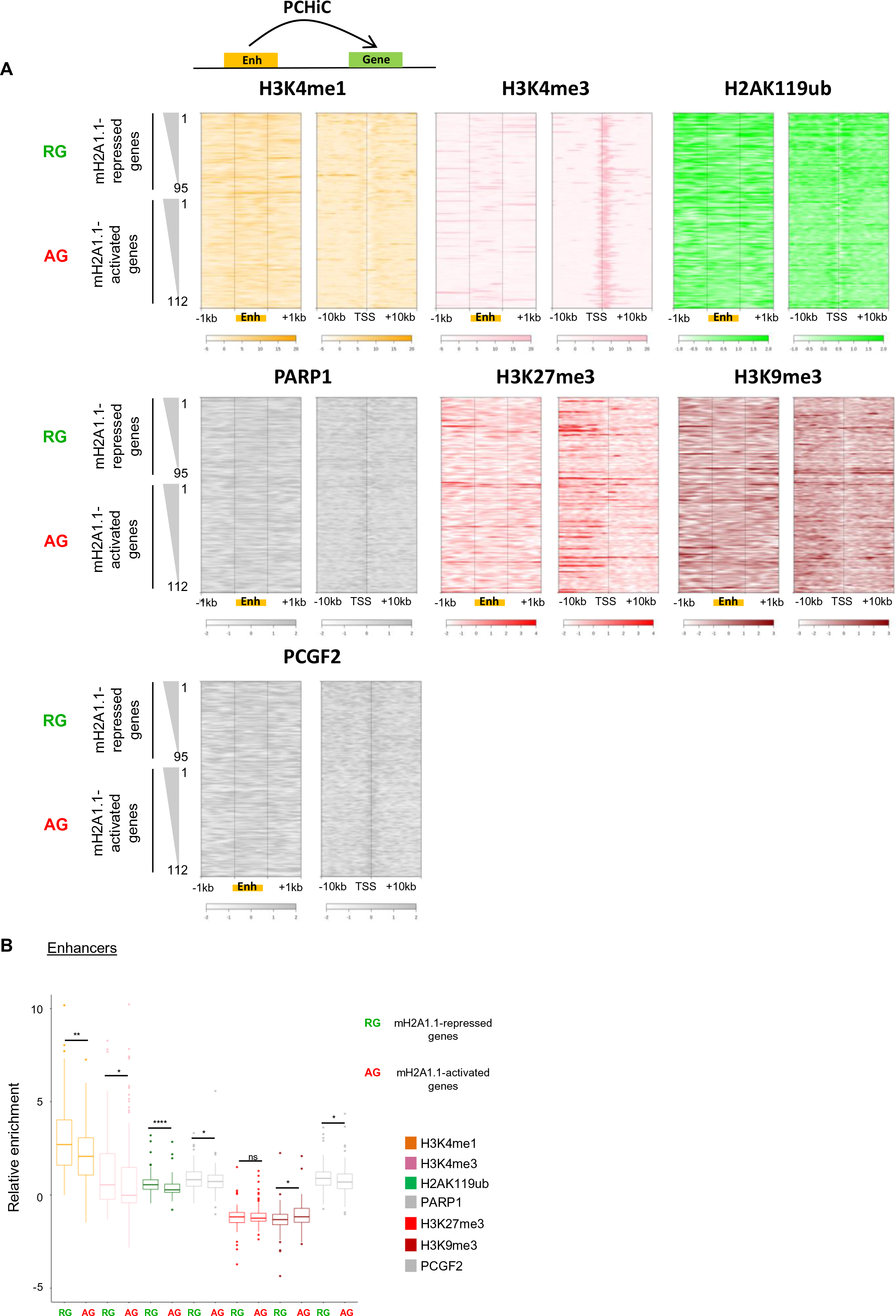
The chromatin environment at enhancers of mH2A1.1-regulated genes. (A) Heatmap profiles showing indicated ChIP-seq data relative enrichment around the TSS (+/- 10 kb) of mH2A1.1-regulated genes (right) and their associated enhancers (+/- 1kb) (left). Same legend as in S12B. (B) Boxplots showing the relative enrichment of indicated ChIP-seq between the enhancers of mH2A1.1-repressed genes and the enhancers of mH2A1.1-activated genes. ns, not significant. “*” : p < 0.05, “**” : p < 0.01, “****” = p-value < 2.2x10^-16^.

**S14 Fig.**
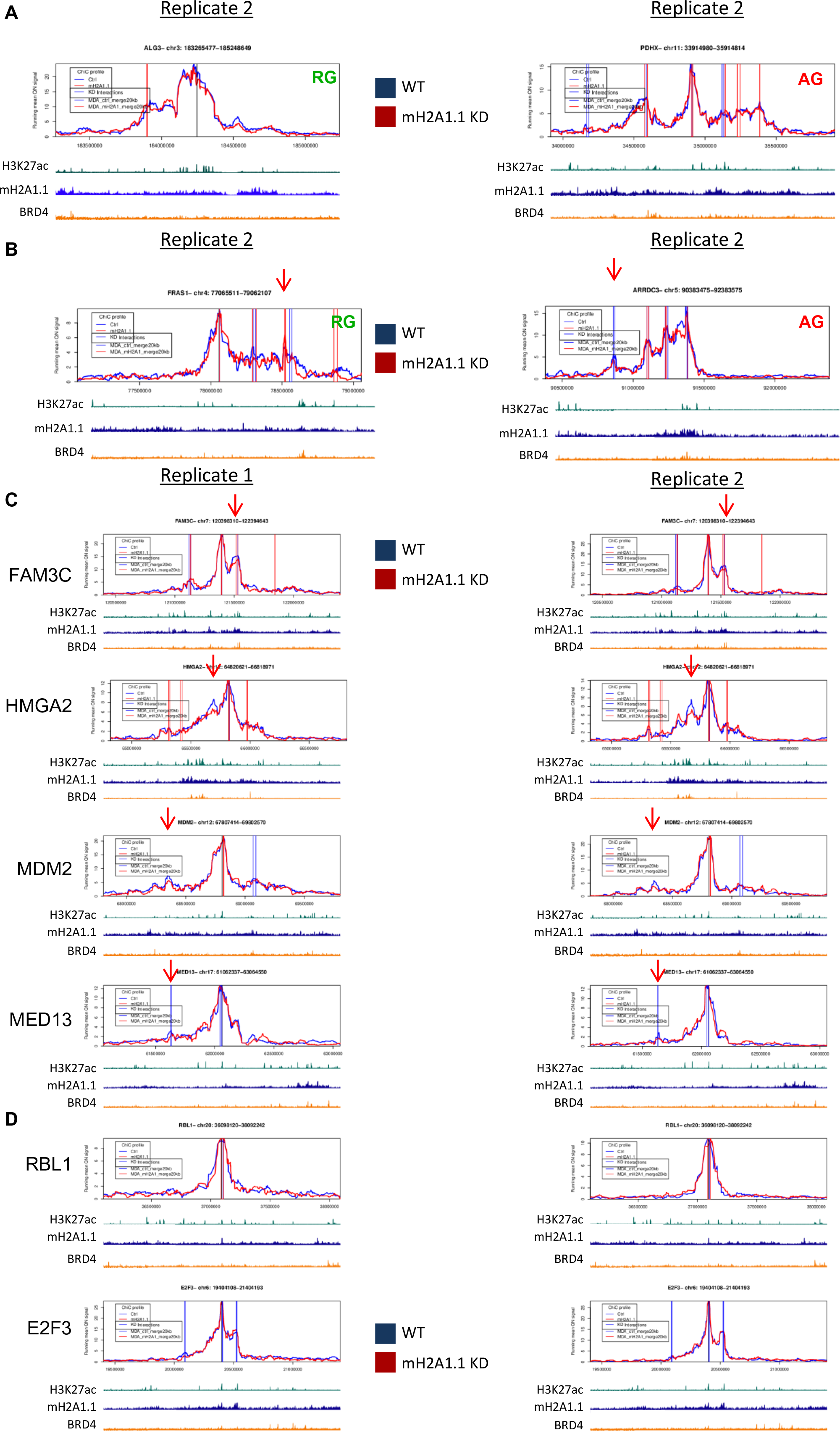
Examples of local genomic interactions of mH2A1.1-target genes. (A) Snapshots of PCHiC data set (replicates n°2) on one mH2A1.1-repressed gene (left) and one mH2A1.1- activated gene (right) in control and mH2A1.1 KD conditions, as indicated. Same legend as in Fig 6C. (B) Same as in (A) but for one mH2A1.1-repressed gene on the top and a mH2A1.1- activated gene on the bottom. Replicates n°1 and 2 are shown. (C) Snapshots of PCHiC data set of 4 mH2A1.1-activated genes as indicated, in control and mH2A1.1 KD conditions. Replicates n°1 and 2 are shown. (D) As in (C) but for two mH2A1.1-activated genes used in Fig 4E. The gene *GTF2H3* was not sequenced in our PCHiC data.

**S15 Fig.**
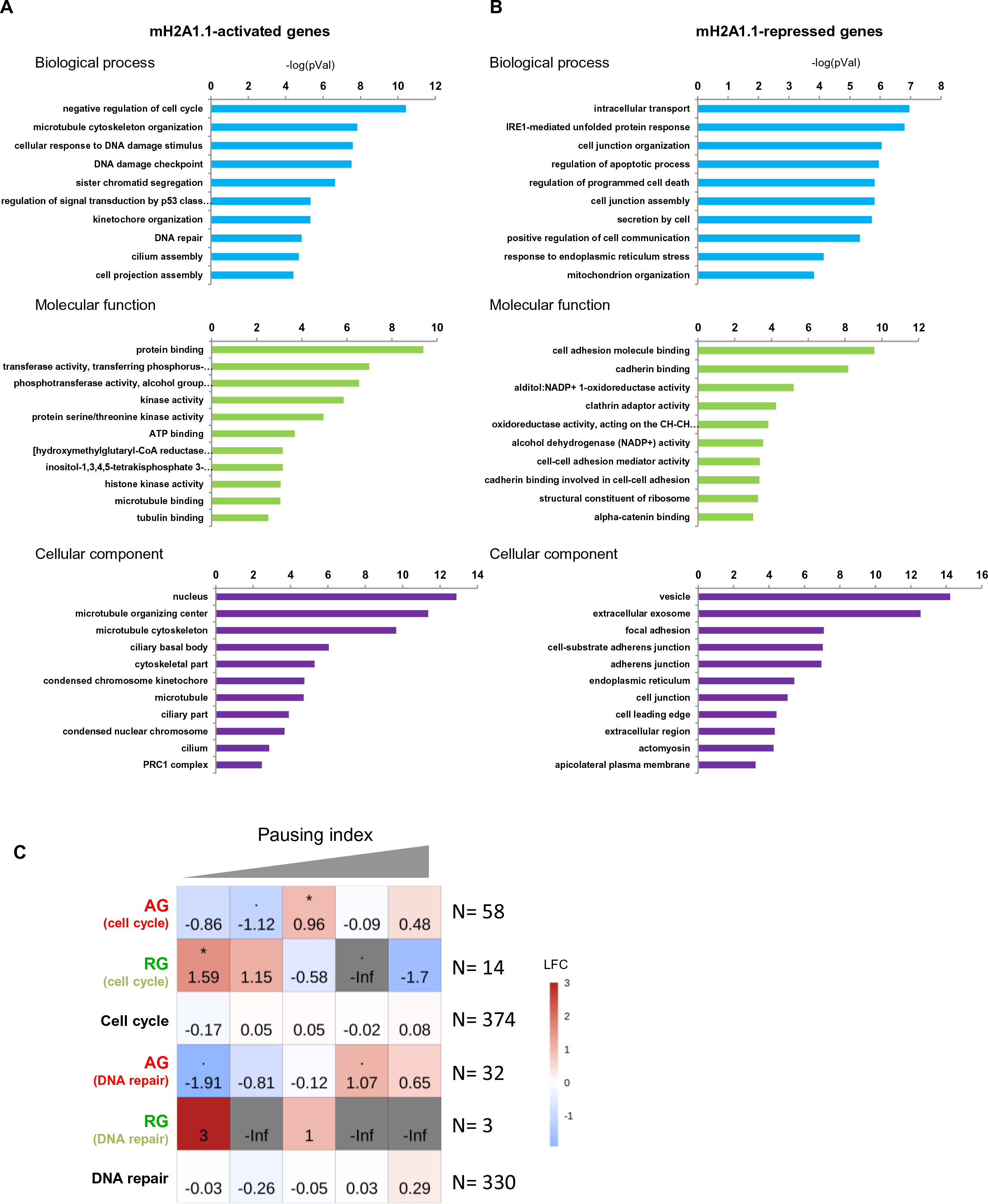
The isoform mH2A1.1 modulates expression of genes involved in cell cycle, DNA repair and cell shape. (A) List of gene ontology (GO) terms for mH2A1.1 activated-genes. The most significantly regulated ontologies were kept, based on their adjusted p-value and are shown in three different classes, Biological Process (upper panel), Molecular function (middle panel) and Cellular Component (lower panel). A full list of enriched GO terms is provided in S4 Table. (B) As in (A) but for mH2A1.1-repressed-genes. (C) Fisher test heatmap showing enrichment of indicated mH2A1.1-target genes with genes divided in 5 equal size categories as a function of their pausing index. Stars indicate the significatively of the fisher exact tests; color map and values present in each scare highlight the log2 odd ratio (LOR) of the fisher exact test. N indicates the number of genes used for the analysis.

**S16 Fig.**
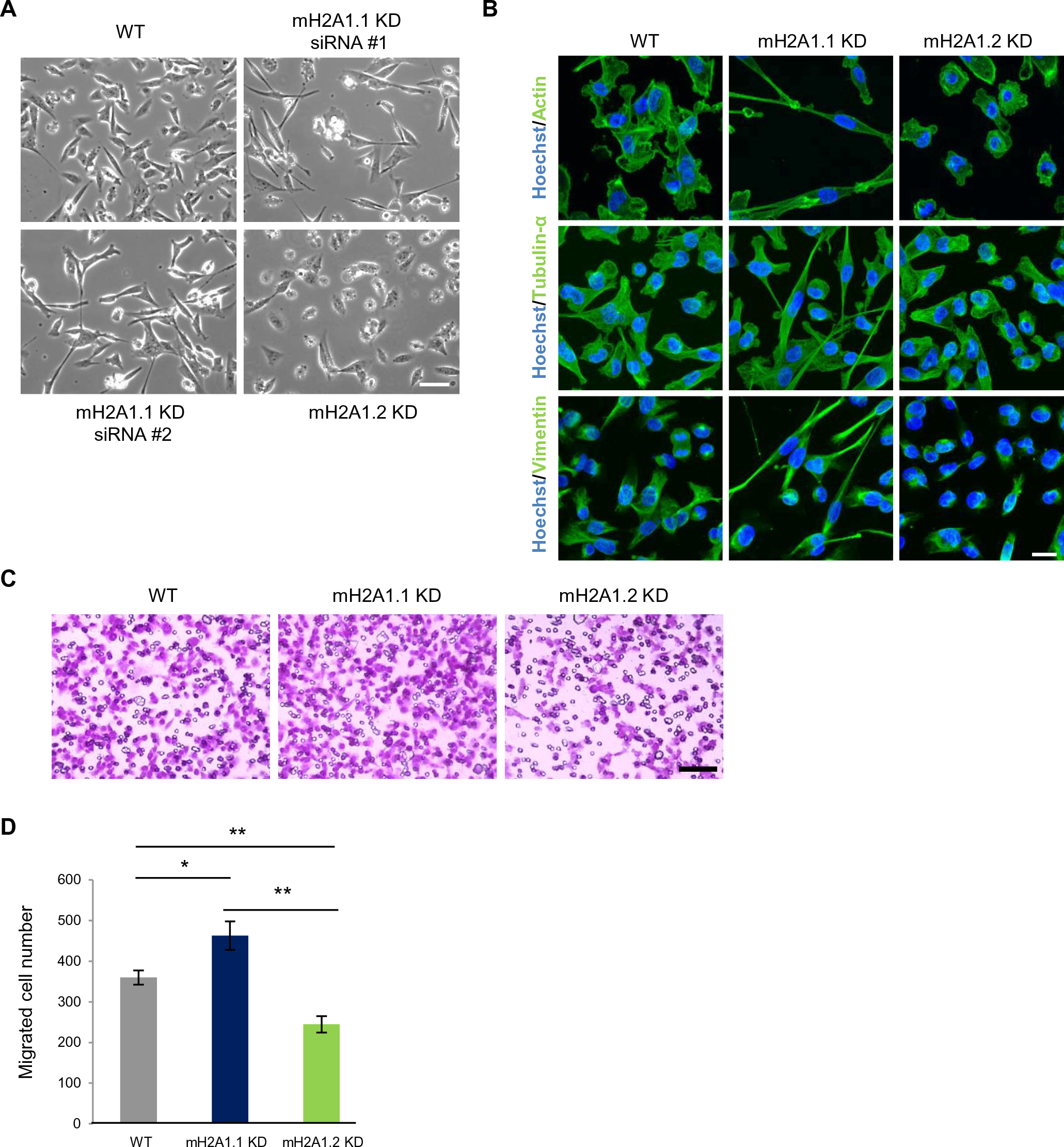
mH2A1.2 promotes cell migration in MDA-MB231 cells. (A) Representative DIC microscopy images of WT, mH2A1.1 KD (two different siRNA) and mH2A1.2 KD MDA-MB231 cells. Scale bar = 100 µM. (B) Immunofluorescence of Actin (up), Tubulin-α (middle) and Vimentin (down) in WT, mH2A1.1 KD and mH2A1.2 KD MDA-MB231 cells. Nuclei are stained with Hoechst. Scale bar = 20 µM. (C) Boyden chamber assay representative images of WT, mH2A1.1 KD and mH2A1.2 KD MDA-MB231 cells. Only migrated cells are labelled in purple. Scale bar = 150 µM. (D) Quantification of Boyden chamber assay presented in (C). Error bar represents s.d from n=3 independent experiments as illustrated in (C). “*” = p-value (p) < 0.05, **, p < 0.01.

## Tables

S1 Table: mH2A1.1-target genes

S2 Table: List of antibodies

S3 Table: List of NGS data

S4 Table: Gene ontology

S5 Table: siRNA sequences

S6 Table: qPCR primers

## Materials and Methods

### Cell culture

MDA-MB231, HEK-293T and MCF7 cell lines were purchased from ATCC, and were maintained and amplified in Dulbecco’s Modified Eagle’s (DMEM) for HEK-93T and MDA-MB231 cells, and in DMEM-F12 for MCF7 cells, supplemented with gentamycin (50 µg/ml) (Gibco), fetal bovine serum (10%, Gibco) and sodium pyruvate (100 mM, Sigma). Cells were maintained in a humidified incubator at 37°C with 5% CO2. Cells lines were regularly tested for mycoplasma infection (MycoAlert, Lonza). In Montpellier, MDA-MB231 cells were cultured in DMEM supplemented with 10% fetal calf serum and penicillin/streptomycin (100 µg/ml each) and regularly tested for mycoplasma infection.

### Transfection of siRNAs and plasmids

At 30-50% confluence, transfection of siRNA (11nM) was performed using INTERFERin (Polyplus-Ozyme) according to the manufacturer’s protocol. Cells in control condition were transfected with INTERFERin without any siRNA. Transfection of plasmid (1μg) was done with FuGene HD (Promega) according to the manufacturer’s protocol. siRNA and plasmids are available in S5 Table. Two- and three-days post plasmid and siRNA transfection respectively, cells were recovered for experiments.

### Western blotting

Cells were lysed and subjected to western blot analysis as previously described (Yang et al., 2014). Briefly, proteins extracts were separated in 10% polyacrylamide (1:125 bisacrylamide:acrylamide) SDS gels, transferred onto nitrocellulose membrane (Bio-Rad) and blocked with PBS-Tween 0.4% - Milk 5% for 1h at RT with rotation. Membranes were then incubated with primary antibodies overnight (O/N) at 4°C in PBS-Tween 0.4% - Milk 5% with rotation (or 1h30 at RT). Primary antibodies are described in the S2 Table. Rabbit anti- mH2A1.1 antibody was generated according to immunization protocol from Agro-Bio - La fierté Saint-Aubin – France. Membranes were next incubated with secondary antibody in PBS-Tween 0.4% - Milk 5% 1h at RT with rotation and the signal was detected using chemiluminescence. Secondary antibodies are described in the S2 Table. Signal quantifications were carried out with Image Lab software (Bio-Rad).

### RNA extraction, reverse transcription and quantitative real-PCR (qRT-PCR)

Total RNA was isolated using the RNAeasy midi kit (Qiagen). Purified RNA was reversed transcribed to cDNA using Maxima H Minus first Strand cDNA synthesis kit (Promega). The sequences of the primers used are available in S6 Table. RT-PCR was performed using iTAq Universal SYBR Green (Bio-Rad) according to manufacturer’s instructions. At least two independent experiments were performed for each condition. The relative expression levels of mRNA were normalized to RPLP0 mRNA expression and evaluated according to the 2^-ΔΔCt^ method (Rao et al., 2013).

### Fluorescence microscopy

Two- or three-days post-transfection, cells were fixed with 4 % paraformaldehyde for 15 min for MDA-MB231 cells and 10 min for HEK-293T at RT. Cells permeabilization was carried out using 0.1 % Triton X-100 in PBS for 10 min at RT. Cells were then blocked with 5 % BSA-0.15% Tween in PBS for 1h at RT. Next, cells were incubated with primary antibody O/N at 4°C. Cells were then incubated with Alexa conjugated secondary antibody for 1h at RT. Actin was labelled using cytoPainter Phalloidin iFluor diluted 1:1000 with secondary antibody according to the manufacturer’s protocol (Abcam, Ab176759). Antibody references and dilutions are provided in S2 Table. The coverslips were finally incubated with Hoechst (Invitrogen, 33342) for 30 min and then mounting with mounting media (Vectashield). Images were acquired with Zeiss LSM 710 big confocal microscope using an x63 PL APO oil DIC On 1.4 objective for all experiments. Images were taken in Z-stacks with a voxel size of 300 nm. A Z-stack or max projection intensity of Z-stacks are shown.

### Chromatin immunoprecipitation and library preparation

Cells were cross-linked in DMEM containing 1.2 % of paraformaldehyde at RT for 10 min with rotation. Cross-link was stopped by the addition of glycine to a final concentration of 0.125M for 5 min. Cell were harvested and lysed in cell lysis buffer (10 mM Tris-HCl pH 7.4, 15 mM NaCl, 60 mM KCL, 1 mM EDTA, 0.1 mM EGTA, 0.2% NP-40, 5% sucrose). After 10 min in ice, cell lysis was amplified with a 2mL dounce (Kimble Chase) to enhance the nuclei separation from cytoplasm. Cell lysis buffer containing lysed cells was deposit up to a pillow buffer (10 mM Tris-HCl pH 7.4, 15 mM NaCl, 60 mM KCL, 1 mM EDTA, 0.1 mM EGTA, 0.2% NP-40, 10% sucrose). Nuclei were then pelleted by centrifugation and wash with washing buffer (10 mM Tris-HCl pH 7.4, 15 mM NaCl, 60 mM KCL). Nuclei were then resuspended in sonication buffer (50 mM Tris-HCl pH 7.5, 150 mM KCl, 5 mM EDTA, 1% NP-40, 0.1% SDS, 0.5 % Sodium deoxycholate, Protease Inhibitor (Roche)). Chromatin was sheared using a Bioruptor (Diagenode) (30 cycles, 30 sec ON/ 30 sec OFF) in order to obtain chromatin fragments with an average size of 300-500 bp. Quality and size of chromatin fragments was monitored by ethidium-bromide stained agarose gel electrophoresis after DNA purification. Then, 100 μg of DNA was incubated with antibody O/N at 4°C on a rotation wheel. Antibodies are described in the S2 Table. 3 mg of protein A magnetic dynabeads (Sigma) were added for 3h at 4°C on a rotation wheel. Immunoprecipitates were then exposed to serial washes for 5 min each on a rotation wheel at 4°C in the following buffers (two times/buffer): WB_I_ :2 mM EDTA, 20 mM Tris pH 8.1, 1 % Triton 100X, 150 mM NaCl, WB_II_: 2 mM EDTA, 20 mM Tris pH 8.1, 1 % Triton X100, 500 mM NaCl, WB_III_: 1 mM EDTA, 10 mM Tris pH 8.1, 250 mM LiCl, 1 % Sodium deoxycholate, 1 % NP- 40 and WB_IV_: 1 M EDTA, 10 mM Tris pH 8.1. Chromatin was eluted from the magnetic beads with DNA isolation buffer (2% SDS, 0.1 M NaHCO3) for 1h at 65°C under agitation. Extracts were reverse-crosslinked with SDS O/N at 65°C. RNAs were degraded with RNase A and proteins were finally degraded with proteinase K. Same procedure was performed for input (10 μg of DNA). DNA was finally extracted with a phenol-chloroform extraction. Quantity and quality of DNA was tested with a nanodrop (NanoDrop2000, Thermo). Samples were sequenced with the GeT core facility, Toulouse, France (http://get.genotoul.fr). Sequencing was done HiSeq3000-HWI-J00115 according to the manufacturer’s protocol. Same procedure was done for ChIPqPCR. The sequences of the primers used fort the qPCR are available in S6 Table. For western blot analysis, extracts (Input (10% IP), No immunoprecipitated (NoIP) fraction and IP fraction were processed as ChIP extract but not incubated with RNAase A and proteinase K. Extracts were then subjected to western blot analysis as previously described in the western blot paragraph. To compare different extracts, we loaded 2 % of Input, 0.5 % of Input, 0.5 % of NoIP fraction and 20% of IP fraction. Percentages are relative to the DNA quantity used for ChIP.

ChIP-seq of H3K27ac, H3K4me1, H3K4me3, H3K36me3 and Pol II were done essentially as previously described (Tolza et al., 2019)(Bejjani et al., 2021). Briefly, after cell fixation with 1 % of paraformaldehyde at RT for 5 min, cells were incubated in cell lysis buffer (PIPES 5 mM, KCL 85 mM, NP40 0.5%, Na Butyrate 10 mM, protease inhibitors) for 10 min on ice. After mild centrifugation, nuclei were lysed in Nuclei Lysis Buffer (Tris-HCL 50 mM pH 7.5, SDS 0.125%, EDTA 10 mM, Na Butyrate 10 mM, protease inhibitors) at 4°C for 2h and, then, sonicated for 10 cycles at 4°C using BioruptorPico device from Diagenode. For immunoprecipitation of H3K4me1, H3K4me3 and H3K27ac, 150 µl of chromatin (equivalent to 4.10^6^ cells) and 4.5 µg of the corresponding antibodies were used. For Pol II, 850 µl of chromatin (equivalent to 22.10^6^ cells) and 20 µg of the corresponding antibody were used. Each ChIP were sequenced by the MGX genomic platform (Montpellier) using the Hi-seq2500 Illumina sequencer. ChIPqPCR of Pol II were done following the same protocol as for Pol II ChIP-seq. qPCR were done on ChIP samples and Input (1% of DNA used for ChIP). qPCR results are normalized using the signal obtained with the input (% of input). Primers used are given in S6 Table.

### Strand-specific total RNA library preparation

Total RNA was isolated using the RNAeasy midi kit (Qiagen). RNA-seq quality and quantity control were performed using a Nanodrop (NanoDrop2000, Thermo) and BioAnalyser. Library preparation and sequencing was done by GeT core facility, Toulouse, France (http://get.genotoul.fr) with the kit TruSeq Stranded total RNA according to manufacturer’s institutions. Sequencing was done HiSeq3000-HWI-J00115 according to the manufacturer’s protocol.

### ChIP-seq data processing

The quality of the reads was estimated with FastQC (Illumina, 1.0.0). Published ChIP-seq data of H3K9me3 (GSM2258862), H3K27me3 (GSM2258850) and corresponding input (GSM2258864) in MDA-MB231 cells were downloaded from GEODATASETS (https://www.ncbi.nlm.nih.gov/geo/, GEO accession number : GSE85158) (Franco et al., 2018), and reanalyzed as subsequently described. Published ChIP-seq data of BRD4 (<u>GSM2862187</u>), RING1B (<u>GSM2862179</u>), PCGF2 (<u>GSM2862185</u>) and H2AK119ub (<u>GSM2862181</u>) in MDA-MB231 cells were downloaded from GEODATASETS (https://www.ncbi.nlm.nih.gov/geo/, GEO accession number : GSE107176) (Chan et al., 2018), and reanalyzed as subsequently described. Published ChIP-seq data of H3.3 (GSM3398219), and corresponding input (GSM3398220) in MDA-MB231 cells were downloaded from GEODATASETS (https://www.ncbi.nlm.nih.gov/geo/, GEO accession number : GSE120313) (Ben Zouari et al., 2019) and reanalyzed as subsequently described. Published ChIP-seq data of PARP1 (GSM1517306) in MDA-MB231 cells were downloaded from GEODATASETS (https://www.ncbi.nlm.nih.gov/geo/, GEO accession number : GSE61916) (Nalabothula et al., 2015) and reanalyzed as subsequently described. Sequenced reads were aligned to the human genome Assembly GRCh38 using STAR (2.5.1) algorithm with defaults parameters (Dobin et al., 2013). Details are supplied in S3 Table. Low quality reads were then filtered out using Samtools (Samtools, options –q 10 -view) (Li et al., 2009). Conversion of BAM files to bigWig files was performed with Bamcompare tool (DeepTools utilities v3.1.3) (Ram et al., 2016). Corresponding ChIP-seq data generated from genomic DNA (Input) were used as control for every bigWig files normalization (options: --normalizeUsing RPKM --operation subtract -- binSize 50 bp --smoothLength 150 bp). Peaks were determined with the enrichR function of NormR package (Helmuth J, 2018). NormR parameters were adjusted depending on the bigwig profiles for each ChIPseq data. mH2A1.1-specific peaks were used for all analysis and correspond to the commun peaks between mH2A1.1 and mH2A1 ChIPseq. Number of peaks for each ChIP-seq data are listed in S3 Table. All downstream analyses were mainly performed with R studio. ChIP-seq signal and peaks positions visualization were obtained with IGV (Thorvaldsdóttir et al., 2013). Boxplots were done with ggplot2 (H. Wickham., 2016). Distributions of mH2A1 isoforms and H3K27me3/H3K9me3 common peaks identified at specific genomic features were calculated using ChIPseeker package with default parameters (**Figs 1E and S4D**) (Yu et al., 2015). Statistical analyses are presented in Statistics and Reproducibility paragraph.

### Identification of “putative” enhancers and super-enhancers

All putative enhancers were determined with ROSE utility tools based on H3K27ac signal outside TSS (+/- 2 kb) to avoid TSS bias (Fig 5A-B) (Blinka et al., 2017). TSS annotation is based on TxDb.Hsapiens.UCSC.hg38.knownGene release (n=25,668 annotated genes). Super-enhancers were determined with ROSE utility tools based on H3K27ac signal (options : stitching_distance = 12.5 kb and TSS_exclusion_zone_size : 2500 bp) (**Fig 5D-E**) (Lovén et al., 2013; Whyte et al., 2013).

### Pol II pausing index calculation

Pausing index (PI) was defined as previously (X. Zhang et al., 2017), which is the ratio of Pol II density in the promoter-proximal region ([-30;300] bp centered on TSS) to the Pol II density in the transcribed regions (TSS + 300 bp to TES). Gene annotation is based on TxDb.Hsapiens.UCSC.hg38.knownGene release. Density of Pol II was calculated using the Pol II bigWig file, normalized using --log2 option (DeepTools utilities v3.1.3) (Ram et al., 2016), and the negative values were replaced by zeros. Genes with a width smaller than 1 kb were excluded from the analysis. Moreover, pausing index were not calculated for the genes having a Pol II density lower than 1.2 in the promoter-proximal region and a Pol II density in the transcribed regions lower than 0. Using this threshold, we only calculated pausing index for transcribed genes having a Pol II binding (n=10,564 genes). “Paused” genes were defined as genes that have a PI upper to 2 (n=7,208) (Day et al., 2016).. “Not paused” genes were defined as genes that have a PI lower to 2 (n=3,356).

### Venn diagrams

Intersection of peaks were determined with the function findOverlaps() from GenomicRanges package (Lawrence et al., 2013). To note that for two ChIP-seq peaks intersections, only number of overlaps is counted and not the number of each peaks contained per overlap. This particularity explained why number of peaks changes between venn diagrams for a same ChIP-seq. The area-proportional Venn diagrams were drawn based on images generated by Vennerable package. Enrichment tests associated to Venn diagrams are explained in Statistics and Reproducibility paragraph.

### Correlation heatmaps

Correlation heatmaps using bigWig of indicated ChIP-seq were done with multiBigwigSummary (with or without the options: -bins) and plotCorrelation (option: - spearman correlation heatmap) from DeepTools utilities (3.1.3) (Ram et al., 2016).

### Fisher test heatmaps

Fisher test heatmaps were done using ggplot2 (H. Wickham., 2016). Each scare on the heatmap shows the results of a fisher exact test between the two groups tested. The positive or negative association between the two groups tested is established by the odd ratio, represented by the “score” (log 2 (Odd Ratio) = LOR) and the color scale, that is proportional to the score. Significatively of the overlap is assessed by the p-value, represented by the stars (* ≤ 0.05 and highly significant when ** ≤ 0.01; *** ≤ 0.001; **** ≤0.0001). Groups used for the analysis were divided in equal size according to the ChIP-seq signal.

### Metagene profiles

Metagene analysis profiles were performed with R Seqplot package (Stempor & Ahringer, 2016) using bigWig files (function getPlotSetArray and plotAverage) centered on TSS (+/- 2kb) or from TSS to TES (+/- 2kb). Profiles correspond to the mean value (+/- the standard error). Values upper to 2 standard deviation (sd), considered as outliers, were removed and not used to generate the profiles. Heatmaps profiles were also performed with R Seqplot package using bigwig files (function getPlotSetArray and plotHeatmap) (Stempor & Ahringer, 2016). Some heatmaps profiles were also ranked according to ChIPseq signal, PI index or gene expression log2Fold change. On all heatmaps, colour intensity reflects level of ChIP-seq enrichment. Colour intensity autoscale were always used excepted for heatmaps **S12B** and **S13A** to compared to relative enrichment between mH2A1.1-target genes and their associated enhancers. On profiles and heatmaps, gene directionality was not ignored, meaning that all gene bodies are artificially placed on the right place of the plots.

### BigWig signal quantification

BigWig signals of indicated ChIP-seq data were calculated with R studio based on bigWig files. For all the figures, the sum bigWig file (bin of 50 bp) was calculated on specific genomic regions and normalized by the width of the specific genomic regions.

### RNA-seq analysis

The quality of the reads was estimated with FastQC (Illumina, 1.0.0). The reads were mapped to the human reference genome GRCh38 using the default parameters of STAR (2.5.1) (Dobin et al., 2013). Details are supplied in S3 Table. Low quality reads and duplicates were then filtered out using SAMtools (Samtools, options -q 10 –view ; -rmdup) (Li et al., 2009). Unstranded normalized bigwig files in RPKM were obtained with Bamcompare tool (DeepTools utilities v3.1.3) (options: --normalizeUsing RPKM --operation subtract -- binSize 50 bp). (Ram et al., 2016). Gene counts were performed with htseq-count utilities with default parameters (0.8.0) (Stempor & Ahringer, 2016). FPKM for all genes were calculated with the formula : FPKM = (RCg x 10^6^ ) / (RCpx L) where RCg corresponds to the number of reads mapped to the gene, RCp to the number of reads mapped to all protein-coding genes and L, the Length of the gene in base pairs. FPKM gene counts in control condition were used to classify genes according to their gene expression level in 4 equal size categories (Silent (n=5,625), low expression (n=5,625), middle expression (n=5,625) and high expression (n=5,625)). FPKM gene counts in mH2A1.1 KD condition were also calculated and used to generated the **Fig 1B**. Differential expression analysis was performed with DESeq2 package (Love et al., 2014) with cutoff |FC| > 1.5 and padj < 0.1. Corresponding volcano plot was done with EnhancedVolcano package (Kevin Blighe, Sharmila Rana, 2018) (**Fig 1A**). The mH2A1.1 KD de-regulated genes are listed in S1 Table.

### Promoter Capture HiC and library preparation

PCHIC data were generated on MDA-MB 231 cells in control and mH2A1.1 KD cell using the siRNA#1 (see Methods part “Transfection of siRNA and plasmids and S5 Table). PCHi-C was essentially performed as in (Schoenfelder et al., 2018), using nearly the same promoter library as (Mifsud et al., 2015) (omitting probes from chromosomes 8, 9 and X).

### Promoter Capture HiC analysis

ChiCMaxima_calling was done using the same default parameters and merging of replicate results as in Ben Zouari et al., after processing the fastq files with custom scripts essentially performing the same analysis as HiCUP (Wingett et al., 2015). Comparison of number of called interactions between subgroups were done **Fig 6A**. Intensity of interactions were estimated based on CHi-C read counts for each biological replicate. Only reads > 5 for each biological replicate were kept and interactions between a bait and the other-end closer than 1.5 Mbp. Finally, for each interaction, read counts were quantile normalized using the function “*normalizeBetweenArrays”* from the LIMMA package (Ritchie et al., 2015). Means between biological replicates were used. ChICMaxima Browser were used to generate PCHiC profiles (https://github.com/yousra291987/ChiCMaxima) (Ben Zouari et al., 2019). ChiCMaxima-called and merged interactions wereoverlapped with enhancers using the findOverlaps function from the R GenomicRanges package (Lawrence et al., 2013).More than one enhancer can significantly be in interaction with mH2A1.1-regulated genes. To simplify, only one enhancer per gene was conserved to generate the heatmaps in **S12B** and **S13A**. Some mH2A1.1-target genes are not present in those heatmaps because they do not have PCHIC-called interactions with an enhancer or did not have PCHi-C capture oligonucleotides.

### GO analysis

GO analysis was performed with LIMMA package (--function goana) (3.8) (Ritchie et al., 2015) and corresponding GO terms are supplied in the S4 Table. Selection of genes related to their functions was done with biomaRt package (function getBM()) (Durinck et al., 2005, 2009). We took genes related to: cytoskeleton (GO:0005856), cell adhesion (GO:0007155), Cilium (GO:0005929) and cell junction (GO:0030054) (n=2,509) using attributes= “ensemble gene_id” annotation from biomaRt package. Overlaps of those genes with mH2A1.1-regulated genes were done (mH2A1.1-activated (n=87/533), mH2A1.1- repressed genes (n=71/412). We took genes related to cell cycle (GO:0007049), cell cycle process (GO:0022402), cell division (GO:0051301) and cell growth (GO:0016049). (n=656). Overlaps of those genes with mH2A1.1-regulated genes were done (mH2A1.1-activated (n=64/533), mH2A1.1-repressed genes (n=18/412). We finally took genes related to DNA repair (GO:0006281), cellular response to DNA damage stimulus (GO:0006974) (n=533). Overlaps of those genes with mH2A1.1-regulated genes were done (mH2A1.1-activated (n=37/533), mH2A1.1-repressed genes (n=4/412). For fisher test heatmap with PI, only genes having a PI were used. N indicates the number of genes used for the analysis in Fig 4B, Fig 7D and Fig 15C.

### Transwell migration assay

Transwell migration assays were performed using Transwell plates with 0.8 µm pore polycarbonate membranes (CormingTranswell, Sigma). Three days post siRNA transfection, MDA-MB231 cells were seeded in the upper chamber without FBS and allowed to invade to the reverse side of the chamber under chemoattractant condition with 10% FBS medium in the lower chamber. Following incubation for 16h at 37°C, the cells were fixed with 3.7% formaldehyde for 2 min at RT. Cells permeabilization was carried out with methanol incubation for 20 min at RT. Cells were then stained with Giesma for 15 min at RT. Same final total cell number between conditions was always checked by wide field microscope to avoid proliferation bias for migratory cell comparison. Not migrated cells were finally removed from the upper chamber by using a cotton swab. Migrated cells adhering to the underside of the chamber were photographed using a light microscope at x200 magnification (Invitrogen EVOS Digital Color Fluorescence Microscope). Cell counting was done with ImageJ in ten different fields per condition (Caroline A Schneider, 2012). Three independent experiments were performed for each condition.

### Statistics and reproducibility

All western blot, RTqPCR and Boyden Chamber assay experiments were repeated at least twice as independent biological replicates and results are presented as mean +/- sd. All statistical analyses were done with R. For Western blot, RTqPCR and Boyden Chamber, Wilcoxon tests were used to compare mean values between conditions. p-values were considered as significant when * ≤ 0.05 and highly significant when ** ≤ 0.01; *** ≤ 0.001; **** ≤0.0001. Fisher exact test were used to performed enrichment test of ChIP-seq peaks. Base sets were defined from all the ChIP-seq data or based on TSS annotations.

### Data availability

ChIP-seq,RNA-seq and PCHiC data have been deposited to GEO under accession number GSE140022. Additional data are available upon reasonable request.

## Acknowledgements

We thank M. Buschbeck from JCLR Institute at Barcelona for kindly providing Flag-mH2A1.1 and Flag-mH2A1.2 expression plasmids. ChIP-seq data against mH2A1.1 and mH2A1 isoforms as well as RNA-seq data were performed in collaboration with the GeT core facility, Toulouse, France (http://get.genotoul.fr), and were supported by France Génomique National infrastructure, funded as part of “Investissement d’avenir” program managed by Agence Nationale pour la Recherche (contract ANR-10-INBS-09) and by the Fondation Recherche Medical (DEQ43940 to O.C team including A.H). We are grateful to the Genotoul bioinformatics platform Toulouse Occitanie (Bioinfo Genotoul) for providing help and/or computing and/or storage resources. We thank Marc Piechaszyk and Sylvain Egloff for critical reading of the manuscript. We acknowledge support from the light imaging Toulouse CBI platform. The work was generously funded by the Institut National du Cancer (INCA PL-BIO- 16-269) to KB.

## Author contributions

L.R, A-C.L and K.B conceived this study. F.M validated custom Ab αmH2A1.1 antibody specificity against mH2A1.1. L.R performed ChIP-seq against mH2A1 isoforms and RNA-seq. I.E-J. and F.B performed ChIP-seq against active histone marks. T.S and N.K performed PCHIC data. L.R, A.H and F.R realized bioinformatic analysis of all ChIP-seq and RNA-seq data. T.S, L.R and A.H realized bioinformatic analysis of PCHIC data. Statistical analyses were done by A.H and L.R. L.R performed all other experimental data. L.R, A-C.L, T.S and K.B designed experiments and interpreted results. L.R, A-C.L and K.B wrote the manuscript with input from all other authors. All authors read and corrected the manuscript.

